# Zyxin and LPP differ in their sensing of strained actin filaments at tricellular junctions

**DOI:** 10.64898/2026.09.18.752733

**Authors:** Katherine K. Tjoelker, Tamsuk Paul, Babli Adhikary, Yashar Bashirzadeh, Sahithya S. Iyer, Kristina M. Herman, Leah M. Beel, Lijuan Xing, Gregory A. Voth, Allen P. Liu, Ann L. Miller

## Abstract

The actomyosin cytoskeleton plays important roles in cell-cell adhesion by generating and responding to forces. Actin-binding proteins support actomyosin networks by reinforcing actin filaments, promoting actin remodeling, and transmitting forces to transmembrane adhesion proteins. One family of actin-binding proteins, LIM domain-containing proteins, is recruited to strained actin filaments. Here, we investigate two members of the LIM domain family, Zyxin and Lipoma-Preferred Partner (LPP), using the embryonic epithelium of *Xenopus laevis*, where actin-associated cell-cell junctions connect cells to promote tissue integrity and barrier function. Specifically, we compare Zyxin and LPP’s response to increased tension at tricellular junctions (TCJs), sites of heightened mechanical strain within the tissue. Upon increased tension, Zyxin and LPP mechanoaccumulate to different extents relative to F-actin, suggesting distinct mechanisms. Our results demonstrate that Zyxin’s and LPP’s LIM domain-containing regions (LCRs) are sufficient for mechanoaccumulation as well as responsible for the difference in mechanoaccumulation, while their N-termini play a regulatory role in their mechanosensitive responses. Docking simulations show that a hydrophobic interaction between the LCR and the “crack site” that forms in F-actin filaments under high force may underlie the LCR’s sensing of strained F-actin. Additionally, the docking simulations reveal that previously-identified conserved residues are present at the binding interface. Mutations of conserved residues in Zyxin or LPP reduce the extent of mechanoaccumulation at TCJs, supporting the docking prediction. Together, this study advances our understanding of Zyxin and LPP’s sensing of strained actin at TCJs under mechanical challenge.

**Significance Statement:** The actomyosin cytoskeleton provides structural support for intercellular connections. This is particularly important at high force levels when actin must be reinforced by actin-binding proteins. Two of those proteins, Zyxin and LPP, stabilize strained actomyosin networks. Using frog embryos, we compare the force-dependent recruitment of these proteins at cell-cell junctions where three cells meet. Despite their similar sequences, Zyxin and LPP respond differently to increased tension. We find that the LIM-domain-containing region (LCR) confers this force-sensing behavior for strained actin at cell-cell junctions. The rest of the protein regulates the extent to which the proteins accumulate. Computational modeling suggests that the accumulation difference between Zyxin and LPP may be due to variations in the LCR and strained actin-binding interface.

## Introduction

Epithelial cells form cohesive sheets separating specialized compartments in the body. Cell-cell junctions called adherens junctions (AJs) adhere epithelial cells to one another, while tight junctions (TJs) establish a barrier that selectively regulates tissue permeability. These cell-cell junctions sense and respond to physiological mechanical forces during developmental morphogenesis and organ-specific functions in adult tissues (1–3). Tricellular junctions (TCJs) are sites where three cells converge and feature specialized proteins and mechanisms, compared to bicellular junctions (BCJs), that allow them to respond to mechanical forces (4–7).

To enable dynamic responses to mechanical forces, cell-cell junction proteins interact with the actomyosin cytoskeleton, comprised of actin filaments (F-actin) and Myosin II, to transmit forces to and provide structural support for junctions (8–9). Increased mechanical force from junction-associated actomyosin bundles that terminate at TCJs and other higher-order vertices (e.g., 4-way junctions or rosettes) generates outward forces on these vertices, challenging junction integrity and barrier function at these sites (10–12). Mechanosensitive scaffolding proteins (e.g., α-catenin, Vinculin, Ajuba (another LIM domain-containing protein), Afadin, ZO-1) connect F-actin bundles to transmembrane proteins (e.g., Angulin, Tricellulin, E-cadherin, Nectin, Claudins) at TCJs (13–21) and can reinforce these connections in a tension-dependent manner to maintain adhesion and barrier function (11,16,22,23) and support processes that involve increasing physiological forces such as cranial neural tube closure (24). The force-sensing mechanisms differ between proteins: Vinculin and Jub/Ajuba respond to a mechanosensitive conformational change in α-catenin (13,14,15), whereas Canoe/Afadin exhibits edge-to-vertex flow in response to mechanical tension that is gated by MBT/PAK in a kinase-independent manner (25) as well as tension-sensitive Abl kinase phosphorylation of Canoe/Afadin that modulates its mechanosensitivity (16). However, it remains unclear how cells sense and respond to changes in F-actin at TCJs under increased mechanical force.

LIM domain-containing proteins are a family of approximately 70 proteins, many of which are mechanosensitive and localize to actin at stress fibers, focal adhesions, and cell-cell junctions in response to mechanical signals (26,27). LIM domain-containing proteins act as scaffolds orchestrating the assembly of protein complexes for a wide range of cellular functions (28). The Zyxin family of LIM domain-containing proteins is composed of seven proteins, including Zyxin and LPP, that bind to strained actin filaments and act as scaffolds to recruit additional proteins to these sites. Mechanical stress results in the phosphorylation of a serine that disrupts the head-tail interaction of Zyxin, allowing the LIM domain-containing region (LCR), made up of three tandem LIM domains, to sense the strained actin at focal adhesions, where it stabilizes and reinforces F-actin through interactions with strained actin filaments (29–33). Molecular dynamics simulations suggest that the LIM domain senses the strained F-actin through binding to a “crack site” that forms in the F-actin filament when it experiences high levels of force (33). The N-terminus of Zyxin then recruits α-actinin, an actin-bundling protein, and VASP, a protein that helps polymerize actin, to repair damaged actin stress fibers (34–36).

While Zyxin is best characterized at stress fibers and focal adhesions, growing evidence also places Zyxin family members at apical cell-cell junctions and junctional vertices, suggesting a functional role at these sites. Zyxin localizes in a tension-dependent manner to apical cell-cell junctions in *C. elegans*, which simultaneously perform the roles of both TJs and AJs (37,38). Our group has also recently reported localization of Zyxin and LPP at apical cell-cell junctions in the embryonic epithelium of *Xenopus laevis* (39). In cultured epithelia, Zyxin and LPP are enriched at cell vertices in MDCK cells (40), and Zyxin accumulates adjacent to apical vertices in the follicular epithelium of *Drosophila* ovaries, consistent with TCJs at this stage of development being under high tension (23). Because Zyxin and LPP can stabilize and recruit factors to repair strained F-actin and are present at TCJs, we hypothesize that Zyxin and LPP sense strained actin within the F-actin bundles that terminate at TCJs to help reinforce these connections.

In this study, we set out to test how Zyxin and LPP respond to mechanical forces in the developing vertebrate epithelium and to elucidate the mechanisms underlying their responses. Using blastula- and gastrula-stage *Xenopus laevis* embryos as an epithelial model, we imaged fluorescently tagged Zyxin and LPP with confocal and super-resolution microscopy. We measured Zyxin and LPP mechanoaccumulation in response to changes in mechanical forces due to embryonic development, cleavage furrow ingression during cell division, hypoosmotic conditions, and extracellular ATP addition. We found that in response to increased mechanical force, Zyxin and LPP recruitment increases at TCJs, with Zyxin accumulating to a greater extent than LPP. The isolated LCRs of both proteins accumulate to a similar extent as the full-length protein, while the N-termini play a regulatory role. Docking simulations provide a mechanistic understanding of the differences in Zyxin and LPP’s sensing of strained actin; the simulations suggest that Zyxin and LPP’s LCRs have different binding strengths at the crack site in strained F-actin and identify residues that may be important at the binding interface. Together, these results establish Zyxin and LPP as mechanistically distinct sensors of strained actin filaments at TCJs, revealing new insights into how epithelial tissues tune vertex reinforcement under mechanical force.

## Results

### Zyxin and LPP are enriched at TCJs in response to physiological increases in force

To examine Zyxin and LPP under physiological changes in tension, super-resolution confocal imaging was performed in live *Xenopus laevis* embryos expressing Zyxin-mNeonGreen or LPP-mNeonGreen. Zyxin and LPP are enriched at TCJs and higher-order vertices compared to BCJs in blastula-stage epithelial tissues (**Fig. 1 *A***).

**Figure 1.**
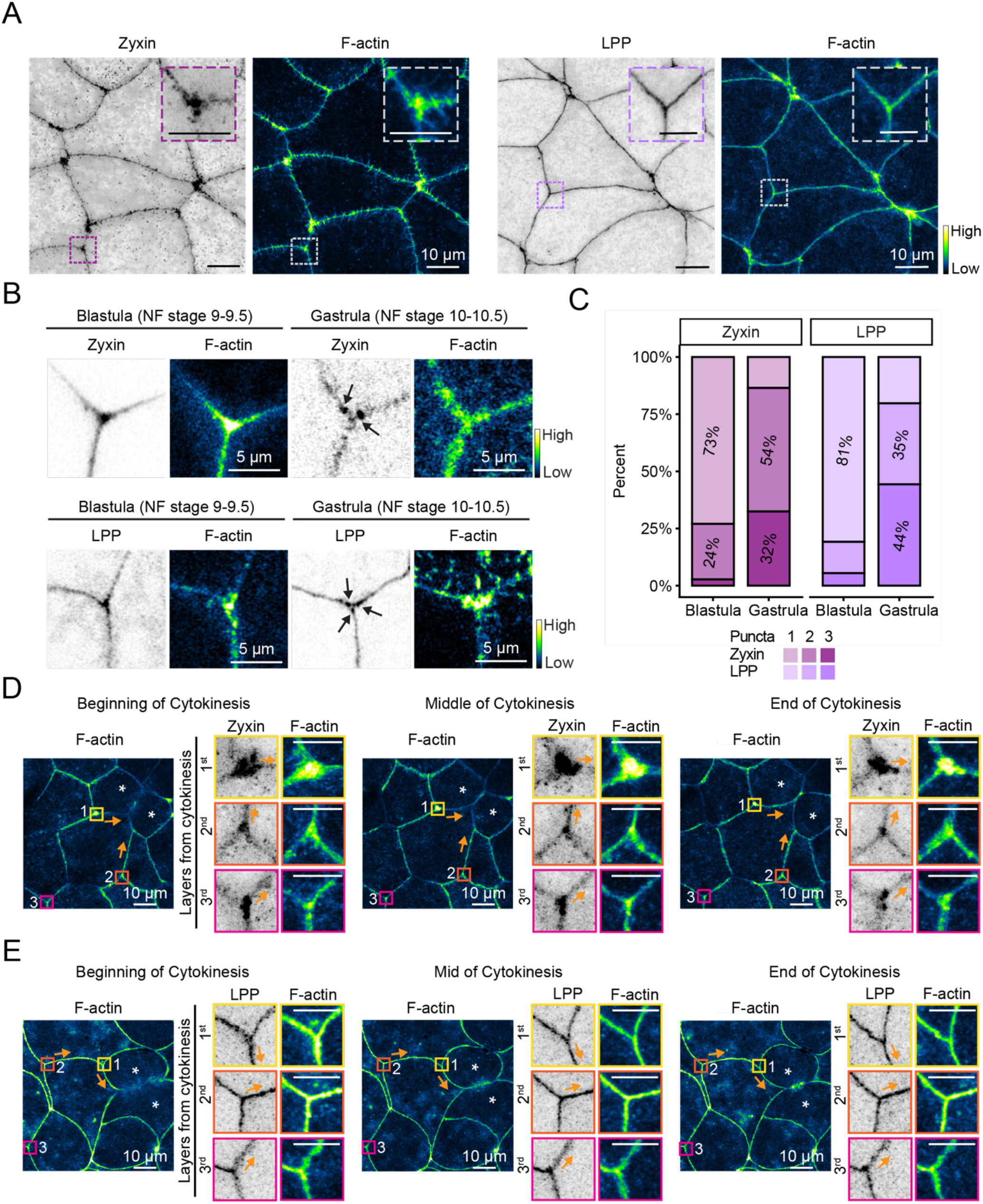
Zyxin and LPP are enriched at and near TCJs, and their spatial organization is tension dependent. *(A)* Live super-resolution images of blastula-stage embryos expressing Zyxin (Zyxin-mNeonGreen, inverted grayscale) or LPP (LPP-mNeonGreen, inverted grayscale) and probe for F-actin (LifeAct-miRFP703, GreenFireBlue lookup table (LUT)). *(B)* Live super-resolution images of blastula and gastrula-stage embryos expressing Zyxin or LPP and F-actin probe. *(C)* Quantification of the number of Zyxin and LPP puncta at TCJs at the blastula and gastrula stage. For Zyxin in blastula-stage embryos, n = 4 clutches, 17 embryos, 74 TCJs; Zyxin in gastrula-stage embryos, n = 4, 11, 37; LPP in blastula-stage embryos, n = 3, 21, 73; LPP in gastrula-stage embryos, n = 4, 19, 79. *(D-E)* Live super-resolution images of cells expressing Zyxin *(D)* or LPP *(E)* and F-actin probe in dividing cells and cells neighboring or near a dividing cell. Scale bar for *(A)*, *(D)*, and *(E)*, 10 µm. Inset scale bars, 5 µm. Scale bar for *(B)*, 5 µm.

Mechanical properties change throughout the early stages of embryonic development (41) and when cytokinesis occurs in a nearby cell (42–43). Embryo stiffness increases as the embryo undergoes gastrulation (41), and tension changes across the animal cap epithelium as it thins and spreads during epiboly (44). Consistent with a developmental change in tissue mechanics, Zyxin and LPP organization at TCJs shifts from a single punctum at late blastula stage (Nieuwkoop and Faber (NF) stage 9-9.5) towards multiple puncta at gastrula stage (NF stage 10-10.5) (**Fig. 1 *B* and *C***). When a cell undergoes cytokinesis, the forces generated by the ingressing cytokinetic contractile ring pull on neighboring junctions (43). We reasoned that as the distance from the cytokinetic furrow increases, tension decreases. Indeed, Zyxin and LPP appear more enriched at the TCJs in the first layer nearest the cytokinetic furrow than at the more distant layers from the furrow (**Fig. 1 *D* and *E***). Together, these data suggest that Zyxin and LPP reorganize at TCJs in response to physiological changes in force, indicating that they likely sense strained actin at or near TCJs.

### Zyxin and LPP mechanoaccumulate at TCJs in response to acute increases in force

To further examine Zyxin and LPP’s response to acute mechanical forces in relation to F-actin, we applied hypoosmotic pressure to increase cell tension (45) or added extracellular ATP to stimulate medial apical actomyosin contraction (22, 46–49) (***SI Appendix*, Fig. S1 *A***). During live imaging, we added H_2_O and extracellular ATP, either separately or together, and monitored F-actin, Zyxin, and LPP enrichment and organization at TCJs pre- and post-addition. In each of these conditions, F-actin intensity increased at BCJs and multi-cellular junctions (MCJs), defined here as TCJs or higher-order vertices, with the combination of hypoosmotic conditions and extracellular ATP addition having the strongest effect, thus demonstrating F-actin’s mechanosensitive response to these mechanical perturbations (**Fig. 2 *A* and *SI Appendix*, Fig. S1 *B-F***). In subsequent experiments, we combined hypoosmotic conditions with the addition of extracellular ATP to characterize the responses of F-actin, Zyxin, and LPP to increased mechanical tension. Hereafter, this condition will be referred to as “extracellular ATP addition”.

**Figure 2.**
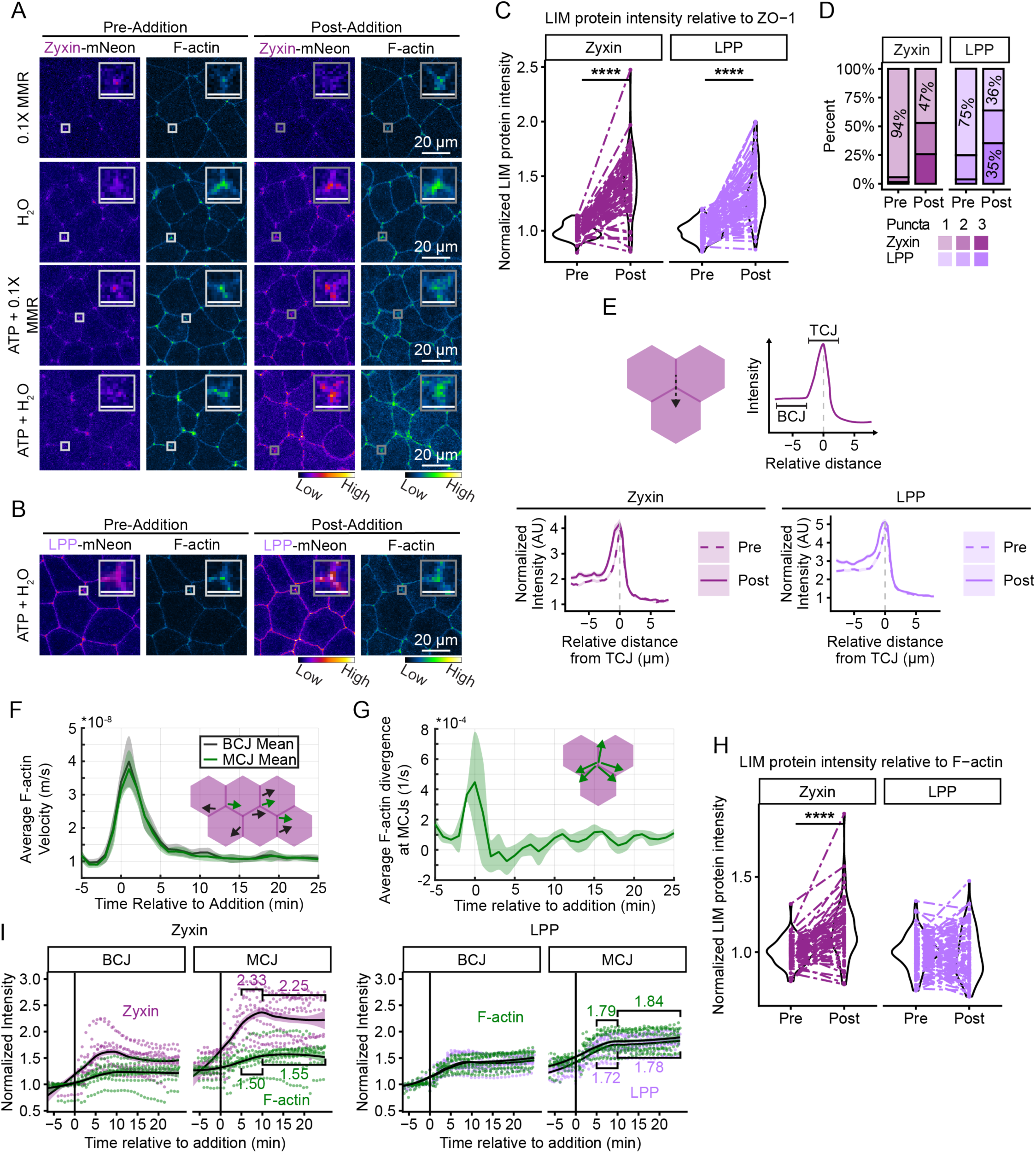
Zyxin and LPP mechanoaccumulate following extracellular ATP addition. *(A)* Live confocal images of cells expressing Zyxin (Zyxin-mNeonGreen, FIRE LUT) and F-actin probe (LifeAct-miRFP703, GreenFireBlue LUT) pre- and post-H_2_O and/or extracellular ATP addition. Enlarged insets highlight changes in Zyxin, and F-actin at TCJs. *(B)* Live confocal images of cells expressing LPP (LPP-mNeonGreen) and F-actin probe (LifeAct-miRFP703) pre- and post-H_2_O and extracellular ATP addition. Enlarged insets highlight changes in LPP, and F-actin at TCJs. *(C)* Quantification of Zyxin and LPP intensity (normalized to ZO-1 intensity) at TCJs pre-and post-extracellular ATP addition. *(D)* Quantification of the number of Zyxin and LPP puncta at TCJs pre- and post-extracellular ATP addition. *(E)* Quantification of normalized Zyxin and LPP signal for linescans that start along a BCJ and then pass through a TCJ and into the cytoplasm of the next cell (see schematic) pre- and post-extracellular ATP addition. *(F)* Quantification of the average velocity of F-actin for BCJs and MCJs. The schematic depicts F-actin velocity vectors determined by PIV. *(G)* Quantification of average divergence of F-actin at MCJs determined by PIV. The schematic depicts F-actin divergence at MCJs. *(H)* Quantification of Zyxin and LPP intensity (normalized to F-actin intensity) at TCJs pre- and post-extracellular ATP addition. *(I)* Quantification of Zyxin, LPP, and F-actin probe intensity at BCJs and MCJs pre- and post-extracellular ATP addition. Data from individual videos are shown as points with LOESS regression function shown as a line. For Zyxin in *(C)*, *(H)*, and *(I)*, n = 3 clutches, 11 embryos, 77 junctions; LPP in *(C)*, *(H)*, and *(I)*, n = 3, 15, 105. Paired t tests in *(C)* and *(H)*: ****p ≤ 0.0001. Scale bar for *(A)* and *(B)*, 20 µm. Inset scale bar, 5 µm.

Extracellular ATP addition reliably led to the mechanoaccumulation of Zyxin and LPP at TCJs relative to the TJ protein ZO-1 or a membrane marker (**Fig. 2 *A*-*C* and *SI Appendix*, Fig. S2 *A and B***). Relative to ZO-1, Zyxin intensity increased by 43 ± 3%, and LPP intensity increased by 35 ± 3%. Additionally, Zyxin and LPP changed their organization at TCJs following extracellular ATP addition from a single punctum to multiple puncta (**Fig. 2 *D***). Furthermore, Zyxin and LPP demonstrated enrichment at TCJs relative to BCJs, and the intensity distribution for Zyxin and LPP moved further away from the TCJ following extracellular ATP addition (**Fig. 2 *E***), consistent with the increased puncta count.

### Velocity of TCJ-associated F-actin increases under mechanical force

Using particle image velocimetry (PIV), we examined the dynamics of junctional F-actin under increased tension. PIV tracking of the actin marker LifeAct has previously been used to characterize actin flow (50). In this study, we use PIV to probe actin flow at cell-cell junctions. This work builds on our finding that Zyxin and LPP localize to regions of stable F-actin at baseline tension (39). Following the addition of extracellular ATP (time 0), F-actin velocity near cell-cell junctions increased for around five minutes, then plateaued near the starting velocity (**Fig. 2 *F***). In contrast, F-actin’s fluorescence intensity increases following ATP addition, then plateaus at the increased level of fluorescence (***SI Appendix*, Fig. S1 *F***). Together, these data suggest a rapid actin response to tension (change in F-actin velocity), then a stable increase in F-actin (F-actin intensity) for at least 25 minutes following the addition of extracellular ATP.

For MCJs specifically, F-actin divergence, which measures the local expansion (sources) or contraction (sinks) of cellular actin, was calculated. In the first few minutes following extracellular ATP addition (time 0), divergence drops below zero (**Fig. 2 *G***). Negative divergence occurs when there is flow into the MCJs. Therefore, this result suggests that acute mechanical force induced by extracellular ATP addition initially leads to F-actin compaction at MCJs.

Following this compaction, divergence rises above 0 indicating an expansion of F-actin at MCJs consistent with previously reported expansion of Myosin II following extracellular ATP addition (22). Together with the velocity data, this demonstrates that F-actin is dynamically reorganized at MCJs in response to mechanical perturbation.

### Zyxin mechanoaccumulates at TCJs and MCJs in a distinct manner from LPP

When Zyxin or LPP intensity is normalized to F-actin, which is also mechanoaccumulating at TCJs **(*SI Appendix*, Fig. S1 *C-F***), Zyxin’s mechanoaccumulation relative to F-actin is significantly increased, but interestingly, LPP does not mechanoaccumulate relative to F-actin (**Fig. 2 *H***). This result demonstrates that Zyxin mechanoaccumulates more than F-actin, whereas LPP mechanoaccumulates to a similar extent as F-actin. Not only do Zyxin and LPP differ in the extent of mechanoaccumulation at TCJs relative to F-actin, but they also differ in their kinetics. Zyxin intensity tends to peak before plateauing at a higher level compared to baseline (**Fig. 2 *I***). Of note, Zyxin intensity peaks at MCJs in a similar timeframe to the F-actin velocity peak (**Fig. 2 *F***). In contrast, LPP shows a small increase in intensity that mirrors the F-actin intensity increase. The odds of a peak in MCJ intensity before the intensity plateaus are 4.2 times higher for Zyxin than for LPP (Pearson’s Chi-squared test p = 0.098). Together, our data suggest that Zyxin senses the increased mechanical force acting on F-actin at TCJs in a distinct manner from LPP.

### Proper pairing between Zyxin and LPP’s N-terminus and LCR is needed for optimal response to mechanical force

To elucidate the differences in mechanoaccumulation between Zyxin and LPP, we generated chimeras that contained the N-terminus from one of the proteins and the LCR from the other (**Fig. 3 *A***). AlphaFold modeling suggests that the first fifty residues in each protein may interact with the LCR (***SI Appendix*, Fig. S3 *A*-*H***). This is consistent with previous work in MDCK cells where the human Zyxin and LPP proteins were shown to have phosphoregulated head-tail interactions (31) and underscores our interest in testing how the proteins behave with a swapped N-terminus/LCR pairing. We imaged the chimeras before and after the addition of extracellular ATP. Prior to the addition of extracellular ATP, the chimeras demonstrated enrichment at TCJs relative to BCJs, as did the wild-type proteins (**Fig. 3 *B* and *C***). Under increased tension, there was an increase in intensity of both fluorescently tagged Zyxin-LPPLCR and LPP-ZyxinLCR (**Fig. 3 *B***). Zyxin-LPPLCR intensity increased by 22 ± 2% and LPP-ZyxinLCR intensity increased by 26 ± 2% (**Fig. 3 *D***). Additionally, the chimeras showed a change in organization from a single punctum to multiple puncta, with LPP-ZyxinLCR showing a more pronounced change than Zyxin-LPPLCR (**Fig. 3 *E***). While both chimeras mechanoaccumulated at TCJs, the extent to which they mechanoaccumulated was reduced compared to the wild-type proteins (**Fig. 3 *F***). This data suggests that the N-terminus/LCR pairing is important for Zyxin and LPP’s optimal response to tension.

**Figure 3.**
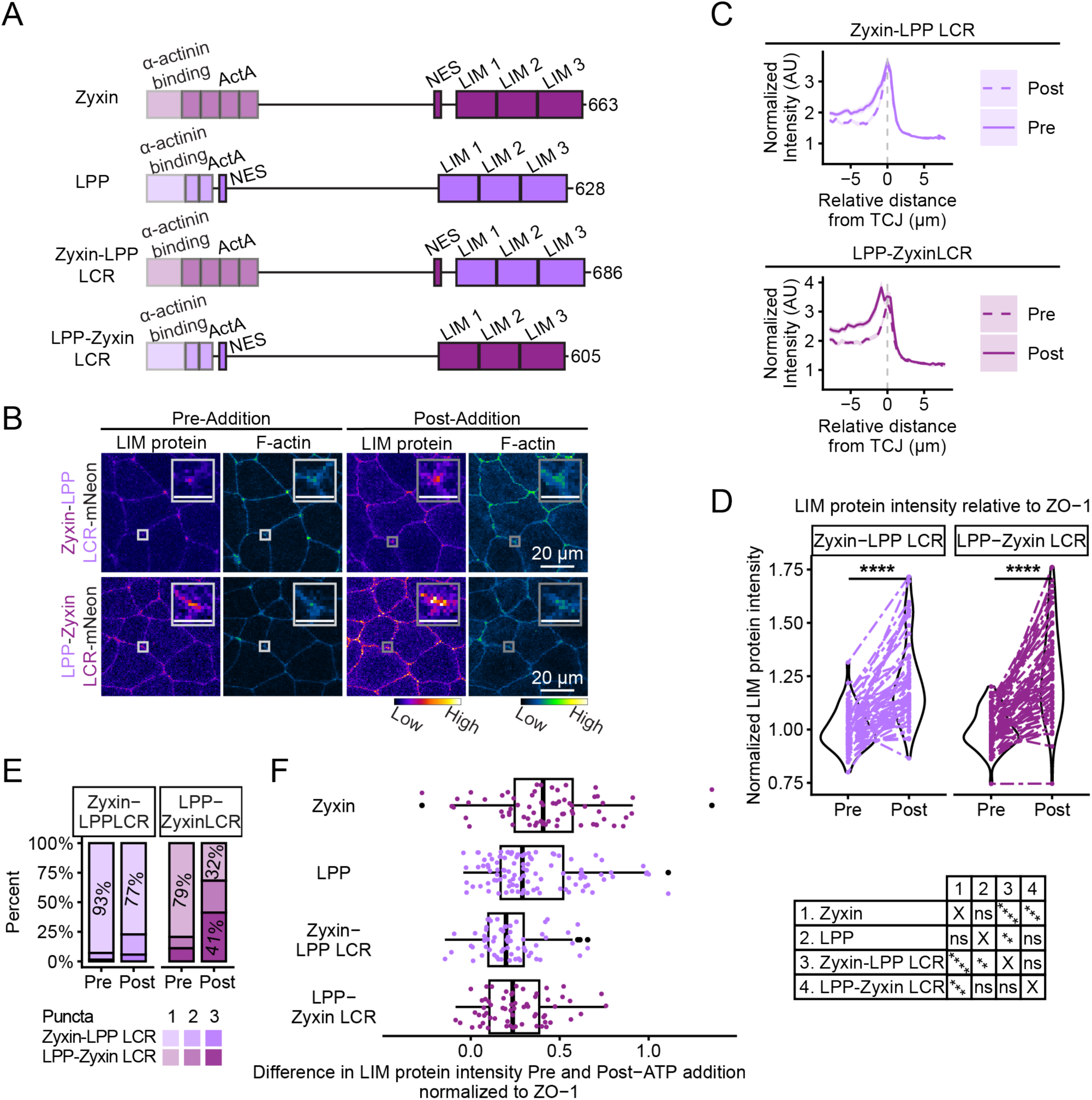
Swapping Zyxin and LPP’s N-terminus and LCR reduces the extent to which the proteins mechanoaccumulate. *(A)* Schematic of wildtype proteins and chimeras in which the LCR of one of the proteins is replaced by the LCR of the other protein. *(B)* Live confocal images of cells expressing Zyxin-LPPLCR (Zyxin-LPPLCR-mNeonGreen, FIRE LUT) and LPP-ZyxinLCR (LPP-ZyxinLCR-mNeonGreen, FIRE LUT) and F-actin probe (LifeAct-miRFP703, GreenFireBlue LUT) pre- and post-extracellular ATP addition. The enlarged insets highlight changes in Zyxin-LPPLCR, LPP-ZyxinLCR, and F-actin mechanoaccumulation at TCJs. Scale bar, 20 µm. Inset scale bar, 5 µm. *(C)* Quantification of Zyxin-LPPLCR and LPP-ZyxinLCR for linescans that start along a BCJ and then pass through a TCJ pre- and post-extracellular ATP addition. *(D)* Quantification of Zyxin-LPPLCR and LPP-ZyxinLCR intensity (normalized to ZO-1 intensity) at TCJs pre- and post-extracellular ATP addition. For Zyxin-LPPLCR, n = 4 clutches, 10 embryos, 70 TCJs; LPP-ZyxinLCR, n = 3, 9, 63. Paired t tests: ****p ≤ 0.0001. *(E)* Quantification of the number of Zyxin-LPPLCR and LPP-ZyxinLCR puncta at TCJs pre- and post-extracellular ATP addition. *(F)* Quantification of the difference in LIM protein intensity (normalized to ZO-1 intensity) pre- and post-extracellular ATP addition. Statistics (shown in chart, right) by ANOVA: ****p ≤ 0.0001, ***p ≤ 0.001, **p ≤ 0.01.

### Zyxin and LPP’s LCRs are sufficient for mechanoaccumulation at TCJs, while the N-termini show minimal TCJ-specific mechanoaccumulation

Based on the observation that the pairing between the N-termini and LCR are important for the extent of mechanoaccumulation at TCJs, we next wanted to test how the isolated LCRs respond to increased mechanical force. We injected fluorescently tagged Zyxin LCR or LPP LCR (**Fig. 4 *A***) into embryos and measured their intensity pre- and post-extracellular ATP addition (**Fig. 4 *B***). The LCRs of both proteins showed a more diffuse localization with minimal localization at BCJs or TCJs at baseline tension (**Fig. 4 *B* and *C***). Unlike the full-length proteins, the LCRs localized to both the nucleus and at cell-cell junctions due to the absence of the nuclear export signals (**Fig. 4 *B***). Upon increased mechanical force, the LCRs of Zyxin and LPP mechanoaccumulate at TCJs (**Fig. 4 *B* and *D***). However, compared to the full-length proteins (**Fig. 2 *A* and *B***), the LCR recruitment appears less continuous along BCJs (**Fig. 4 *B***). Notably, unlike the chimeras, the LCRs mechanoaccumulate to a similar extent as the full-length proteins (**Fig. 4 *E***): Zyxin LCR’s intensity increases by 33 ± 4%, and LPP LCR’s increases by 32 ± 2% (**Fig. 4 *D***). Additionally, the Zyxin LCR exhibits a peak in fluorescence 5-10 minutes post-extracellular ATP addition, similar to full-length Zyxin (**Fig. 4 *F***). These data indicate that the Zyxin and LPP LCRs are sufficient for mechanoaccumulation at TCJs.

**Figure 4.**
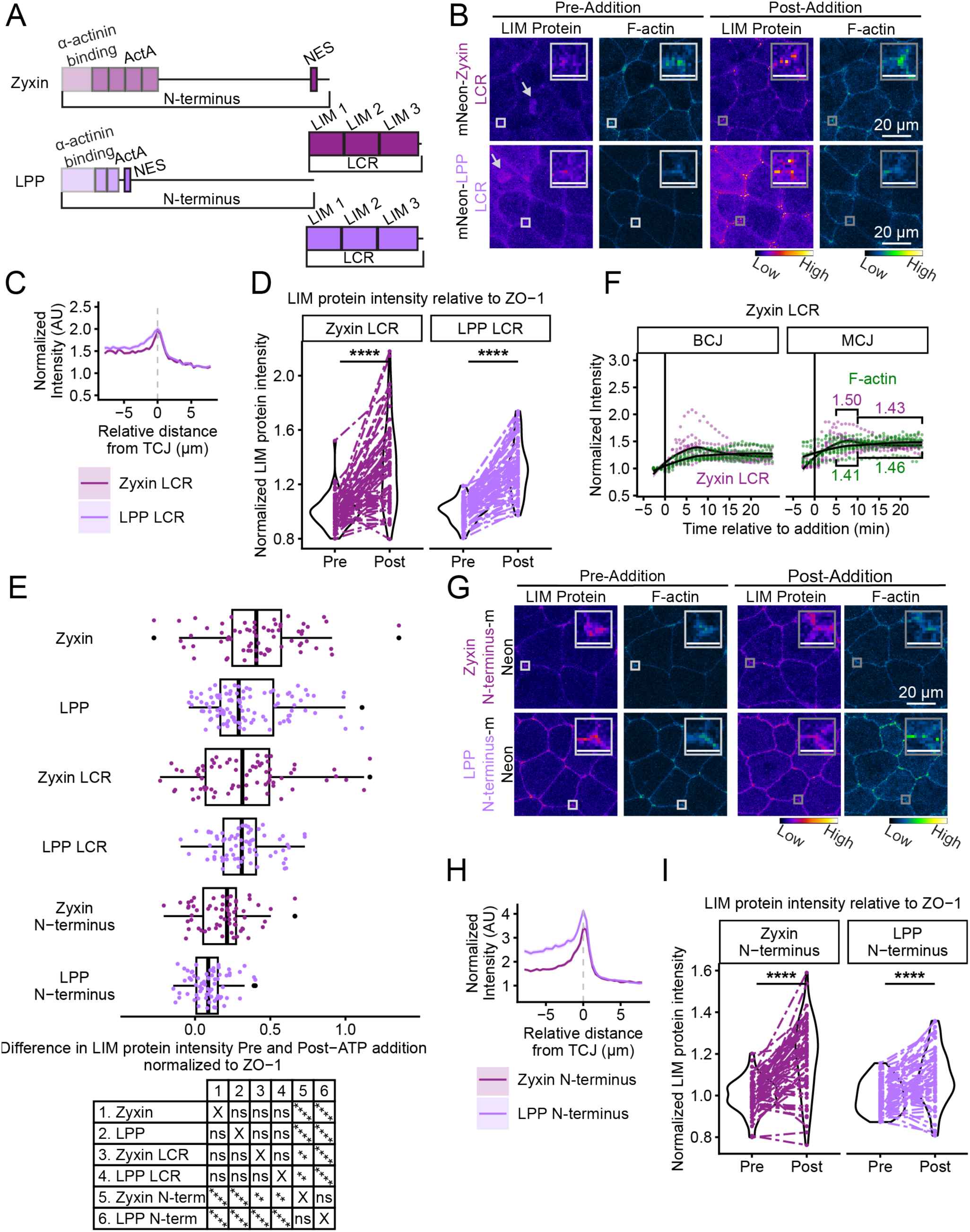
Zyxin and LPP’s LCRs mechanoaccumulate to a similar extent as the wild-type proteins. *(A)* Schematic of Zyxin and LPP N-termini and LCRs. *(B)* Live confocal images of cells expressing Zyxin’s LCR (mNeonGreen-ZyxinLCR, FIRE LUT) or LPP’s LCR (mNeonGreen-LPPLCR, FIRE LUT) and F-actin probe (LifeAct-miRFP703, GreenFireBlue LUT) pre- and post-extracellular ATP addition. The enlarged insets highlight changes in Zyxin LCR, LPP LCR, and F-actin mechanoaccumulation at TCJs. Arrows point to the LCRs’ nuclear localization. *(C)* Quantification of ZyxinLCR and LPPLCR at baseline tension for linescans that start along a BCJ and then pass through a TCJ. *(D)* Quantification of ZyxinLCR and LPPLCR intensity (normalized to ZO-1 intensity) at TCJs pre- and post-extracellular ATP addition. For ZyxinLCR, n = 4 clutches, 10 embryos, 70 TCJs; LPPLCR, n = 3, 10, 70. Paired t tests: ****p ≤ 0.0001. *(E)* Quantification of the difference in LIM protein intensity (normalized to ZO-1 intensity) pre- and post-extracellular ATP addition. Statistics (shown in chart, bottom) by ANOVA: ****p ≤ 0.0001, ***p ≤ 0.001, **p ≤ 0.01. *(F)* Quantification of ZyxinLCR and F-actin probe intensity at BCJs and MCJs pre- and post-extracellular ATP addition. Data from individual videos are shown as points, with the LOESS regression function shown as a line. *(G)* Live confocal images of cells expressing Zyxin’s N-terminus (ZyxinNterm-mNeonGreen, FIRE LUT) or LPP’s N-terminus (LPPNterm-mNeonGreen, FIRE LUT) and F-actin probe (LifeAct-miRFP703) pre- and post-extracellular ATP addition. The enlarged insets highlight changes in ZyxinNterm, LPPNterm, and F-actin mechanoaccumulation at TCJs. *(H)* Quantification of Zyxin N-terminus and LPP N-terminus at baseline tension for linescans that start along a BCJ and then pass through a TCJ. *(I)* Quantification of ZyxinNterm and LPPNterm intensity (normalized to ZO-1 intensity) at TCJs pre- and post-extracellular ATP addition. For Zyxin-LPPLIM, n = 4 clutches, 10 embryos, 70 TCJs; LPPNterm, n = 3, 9, 63. Paired t test: ****p ≤ 0.0001. Scale bar for *(B)* and *(G)*, 20 µm. Inset scale bar, 5 µm.

Since the N-terminus of Zyxin and LPP regulates the LCR’s optimal response to tension, we tested how the isolated N-terminus of Zyxin and LPP responds to increased mechanical force. We injected mRNA of fluorescently tagged Zyxin N-terminus or LPP N-terminus into embryos and measured their intensity pre- and post-extracellular ATP addition (**Fig. 4 *G***). Zyxin’s and LPP’s N-termini showed a reduction in TCJ enrichment compared to the full-length proteins (**Fig. 4 *G* and *H***). The N-termini showed a small but statistically significant increase in intensity at TCJs following extracellular ATP addition. Zyxin N-terminus intensity increased by 18 ± 2%, while LPP N-terminus intensity increased by 9 ± 1% (**Fig. 4 *I***). This level of mechanoaccumulation is significantly smaller than not only the full-length proteins but also the chimeras and the isolated LCR constructs (**Fig. 4 *E***). Taken together, our data suggest that while the N-termini of Zyxin and LPP show some recruitment to cell-cell junctions in response to mechanical force, the LCRs of Zyxin and LPP are the major force-sensing domains and determine the dynamics and extent of accumulation on F-actin associated with TCJs.

### Crack site 1 is the preferred F-actin crack site for tandem LIM domain docking

To examine the mechanism of LCR force sensing at TCJs, we used docking simulations to model Zyxin LCR and LPP LCR binding to cracked F-actin. Zsolnay et al. reported that applying tension to actin filaments *in silico* exposes different cracked actin interfaces, namely crack site 1 (CS1) and crack site 2 (CS2) (33). CS1 is located between subdomain 2 (SD2) of F-actin monomer ‘i’ and SD1 and 3 of monomer ‘i-2’ and exposes residues from the D-loop (V43, M44, and V45) of monomer ‘i’ and the W-loop (Y169) and C-terminal region (I369 and F375) of monomer ‘i-2’. CS2, located at the interface of SD4 of monomer ‘i’ and SD1 of ‘i-2’, exposes residues including E72 from monomer ‘i-1’and S199, R183, and Q246 from monomer ‘i’. The authors also established through their docking simulations that isolated LIM domains dock preferentially to CS1.

In our study, we investigated whether this F-actin crack site preference holds when LIM domains are presented in their physiological context - three tandem LIM domains making up the LCR, in which they can engage cooperatively or competitively. The LCR sequences of *Xenopus* Zyxin or LPP were docked to the actin filament crack sites (**Fig. 5 *A* and *B***, representative examples of LPP are shown) using the information-driven protein-protein docking platform HADDOCK 2.4 (51,52). Post docking, we estimated the binding energies using the Molecular Mechanics-Generalized Born Surface Area (MM-GBSA) approach implemented in HawkDock (53,54). Comparing MM-GBSA binding energies of all the CS1 and CS2 bound complexes revealed that binding to CS1 is more favored than CS2 for almost all the docked systems except for the Zyxin LIM1-bound model (**Fig. 5 *C***).

**Figure 5.**
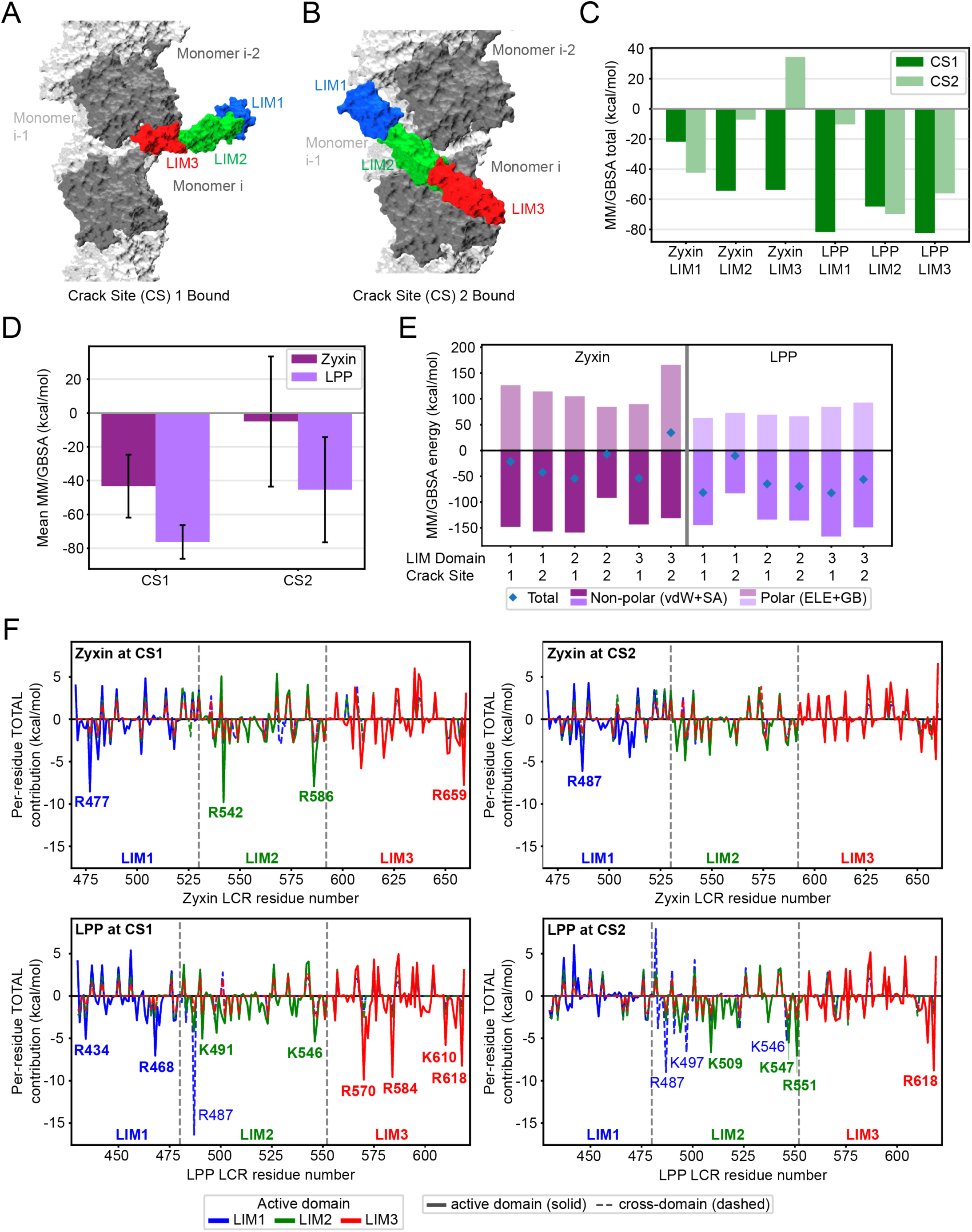
Computational modeling of Zyxin and LPP’s LIM domain docking to F-actin crack sites. *(A-B)* Representative views of *Xenopus* LPP LIM3-bound to crack site 1 (CS1) (*A*) or *Xenopus* LPP LIM2 bound to crack site 2 (CS2) (*B*) docked on a cracked actin filament. LIM1 is blue, LIM2 is green, and LIM3 is red. (*C*) Binding energies of LIM1-3 of *Xenopus* Zyxin and LPP at CS1 and CS2 estimated using MM-GBSA. *(D)* Mean value (across 3 LIM domains) for total binding energy for Zyxin and LPP at CS1 and CS2. Error bars denote standard deviation. *(E)* Total polar and nonpolar energy contributions for LIM1–3 of *Xenopus* Zyxin and LPP bound at CS1 and CS2 across the 12 docking models. *(F)* Residue-wise binding energy decomposition of Zyxin and LPP at CS1 and CS2. The active domain refers to the LIM domain defined as the docking interface in each docking simulation, with contributions from these residues shown as solid lines. Cross-domain contributions arise from residues outside the defined docking interface and are shown as dashed lines.

### LPP exhibits stronger simulated binding to cracked F-actin than Zyxin

To explore the differences in mechanoaccumulation between Zyxin and LPP, we compared the binding energies of Zyxin and LPP’s LCRs to cracked actin. Surprisingly, LPP exhibited more favorable mean MM-GBSA binding energies than Zyxin at both crack sites (**Fig. 5 *D***). At CS1, the mean binding energy was −76.3 ± 10.0 kcal/mol for LPP compared with −43.3 ± 18.6 kcal/mol for Zyxin. Similarly, at CS2, LPP showed a mean binding energy of −45.4 ± 31.1 kcal/mol, whereas Zyxin exhibited a substantially weaker binding energy of −5.0 ± 38.5 kcal/mol. MM-GBSA decomposition suggests that favorable binding of Zyxin and LPP’s LCRs at both F-actin crack sites is dominated primarily by stabilizing non-polar interactions (**Fig. 5 *E***). Notably, the non-polar contributions were very similar between the two proteins: −148.6 ± 16.7 versus - 150.3 ± 8.2 kcal/mol at CS1 and −122.7 ± 34.8 versus −126.7 ± 33.0 kcal/mol at CS2 for LPP and Zyxin, respectively. The difference therefore does not arise from substantially better hydrophobic packing by LPP. Rather, LPP pays a smaller polar penalty than Zyxin at both sites, +72.3 ± 11.0 kcal/mol for LPP versus +107.0 ± 18.4 kcal/mol for Zyxin at CS1 and +77.3 ± 13.9 kcal/mol for LPP versus +121.7 ± 41.2 kcal/mol for Zyxin at CS2 (**Fig. 5 *E***). Physically, the polar term is the sum of two opposing effects - the favorable Coulombic interactions that charged and polar groups form at the F-actin-LIM interface, and the unfavorable energetic cost of desolvating those groups (i.e., removing their bound water as they form the binding interface). A positive value means the desolvation cost outweighs the Coulombic gain, so the polar forces oppose binding overall. This suggests that LPP and Zyxin pack against F-actin similarly but enhanced electrostatic and polar complementarity at the actin-LPP interface contributes to its stronger binding strength.

### Identification of potential actin-binding residues in Zyxin and LPP’s LCRs

Analysis of the total binding energy contribution for all the LCR residues across the 12 combinations (3 LIM domains x 2 crack sites x 2 proteins) identifies recurrent residues in Zyxin and LPP. Positively charged arginine and lysine residues contribute to the binding energy substantially across both the crack sites as well as multiple binding poses (i.e., for different directly engaging LIM domains) (**Fig. 5 *F***). Notably, some residues contribute strongly even when they are not part of the actively engaging LIM domain (cross-domain contributions). For example, in the LPP-LIM1-active pose at CS1, the strongest favorable LCR residue is R487 (−16.37 kcal/mol), which belongs to LIM2 and forms salt-bridge-like contacts with E361 and E364 of actin monomer ‘i-2’. Direct cross-domain engagement occurs for both LCRs but is more extensive for LPP (***SI Appendix*, Table S1**). This result suggests that for both Zyxin and LPP, binding of the tandem LIM domains to actin involves more than one LIM domain to achieve stronger binding. This result is consistent with experimental data showing that removing either LIM1 or LIM3 from Zyxin’s or LPP’s LCRs nearly eliminates mechanoaccumulation at TCJs (***SI Appendix*, Fig. S4 *A*-*D***).

### Role of conserved residues at the LIM domain-F-actin crack site interface

Conserved aromatic residues in each LIM domain are a defining feature of LIM domain-containing proteins that are classified as mechanoresponders (29). These conserved residues were shown to be necessary, though not sufficient, for Zyxin’s mechanoaccumulation at actin stress fibers (29). The residues, which were identified in the human Zyxin and LPP sequences, are conserved aromatic residues (phenylalanine or tyrosine) in *Xenopus* (***SI Appendix*, Fig. S5 *A***). More broadly, the LCR regions of Zyxin and LPP are highly conserved between human and *Xenopus*: Zyxin’s LCR is 75.5% similar to humans, while LPP’s LCR is 95.0% similar to humans (***SI Appendix*, Fig. S5 *B***). When we examined the conserved residues’ role in binding the F-actin crack sites *in silico*, a subset of these residues lay within 5 Å of actin (e.g., Zyxin F642 in the LIM3-active pose at CS1, Zyxin Y512 and F642 in the LIM1- and LIM3-active poses at CS2, respectively, and LPP Y601 in the LIM3-active pose at CS1). The clearest favorable contributions were observed for Zyxin Y512 at CS2, Zyxin F642 at CS2, and LPP Y601 at CS1 (***SI Appendix*, Fig. S5 *C***). These results suggest that the conserved aromatic positions are not the dominant energetic hotspots in the tandem LIM domain docking models. Instead, most of the direct binding energy arises from nearby positively charged arginine/lysine residues and hydrophobic residues (**Fig. 5 *F***), whereas the conserved aromatic positions may be important for the surrounding interaction surface.

### Mutation of conserved aromatic residues in the LCR reduces Zyxin and LPP mechanoaccumulation at TCJs

We then tested the impact of these conserved aromatic residues on mechanoaccumulation at TCJs, given that they may be involved at the interaction surface. We generated alanine point mutations of the conserved residue in each LIM domain (Zyxin-3A and LPP-3A) (**Fig. 6 *A***) and imaged the fluorescently tagged mutants pre- and post-extracellular ATP addition. Interestingly, we found that when the conserved aromatic residues were mutated, Zyxin and LPP retained some ability to mechanoaccumulate at TCJs relative to ZO-1 (**Fig. 6 *B-C***). However, they accumulated to a reduced extent compared with the wild-type proteins: Zyxin-3A and LPP-3A intensity both increased by 17 ± 2% (**Fig. 6 *C* and *D***). Additionally, Zyxin and LPP intensity at TCJs was actually reduced relative to F-actin following extracellular ATP treatment (**Fig. 6 *E***). Regarding the kinetics of mechanoaccumulation, the point mutations of the conserved aromatic residues also eliminated Zyxin’s early intensity peak before the plateau (**Fig. 6 *F* and Fig. 2 *I***). Together, this data demonstrates that while the three conserved aromatic residues contribute to Zyxin and LPP force sensing, there is still some level of mechanoaccumulation at TCJs in the mutants, which is consistent with our computational modeling suggesting that positive residues (arginine/lysine) may have a greater impact on binding strength.

**Figure 6.**
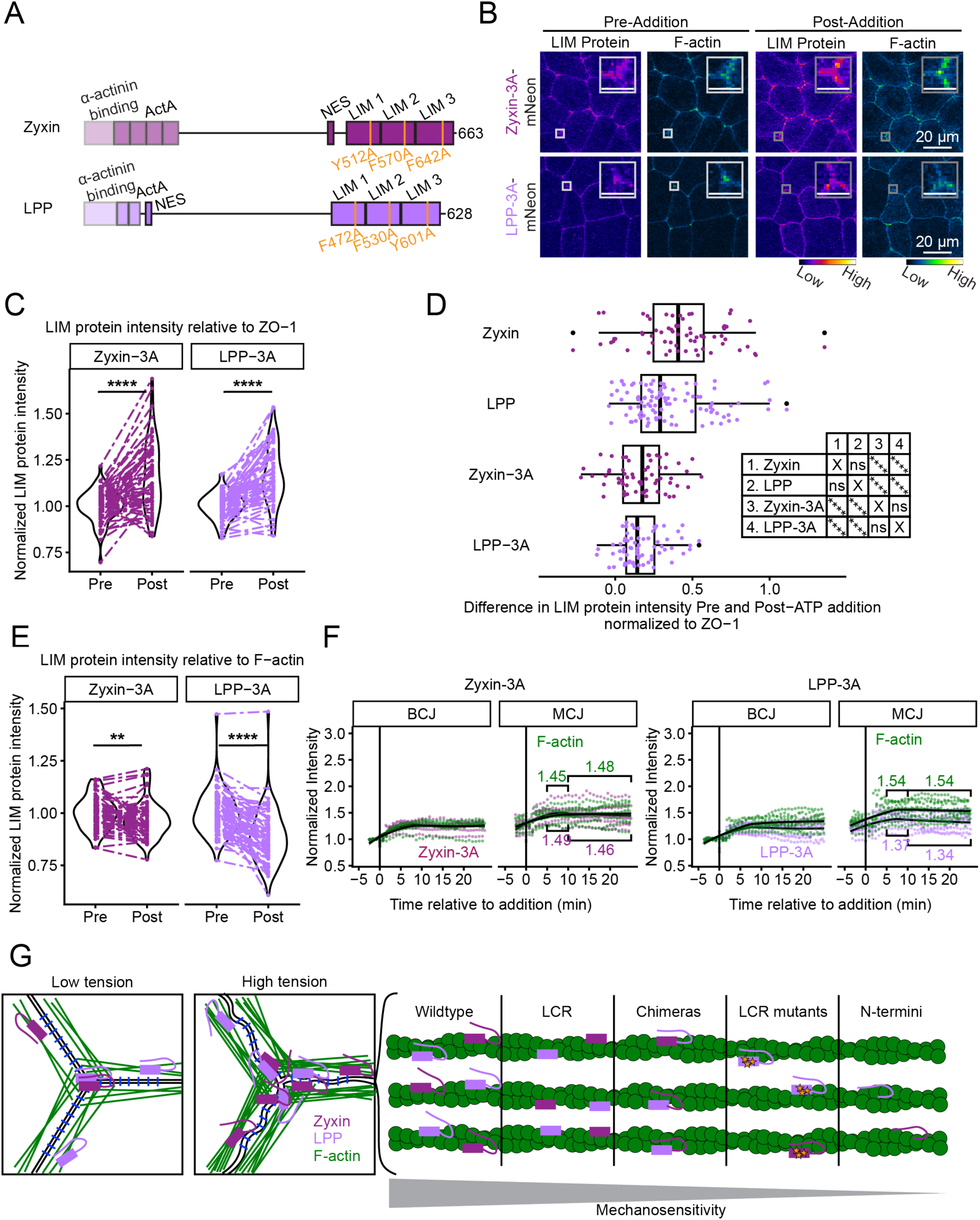
Mutations of conserved aromatic residues in the LCR reduce Zyxin and LPP mechanoaccumulation at TCJs. *(A)* Schematic of Zyxin and LPP with mutated conserved residues labeled. *(B)* Live confocal images of cells expressing Zyxin-3A (Zyxin-3A-mNeonGreen, FIRE LUT) or LPP-3A (LPP-3A-mNeonGreen, FIRE LUT) and F-actin probe (LifeAct-miRFP703, GreenFireBlue LUT) pre- and post-extracellular ATP addition. The enlarged insets highlight changes in Zyxin-3A, and F-actin mechanoaccumulation at TCJs. Scale bar, 20 µm. Inset scale bar, 5 µm. *(C)* Quantification of Zyxin-3A and LPP-3A intensity (normalized to ZO-1 intensity) at TCJs pre- and post-extracellular ATP addition. *(D)* Quantification of the difference in LIM protein intensity (normalized to ZO-1 intensity) pre- and post-extracellular ATP addition. Statistics (shown in chart, right) by ANOVA: ****p ≤ 0.0001. *(E)* Quantification of Zyxin-3A and LPP-3A intensity (normalized to F-actin intensity) at TCJs pre- and post-extracellular ATP addition. For Zyxin-3A in *(D)* and *(E)*, n = 3 clutches, 9 embryos, 63 TCJs; LPP-3A in *(D)* and *(E)*, n = 6, 11, 77. Paired t tests in *(D)* and *(E)*: ****p ≤ 0.0001, **p ≤ 0.01, *p ≤ 0.05. *(F)* Quantification of Zyxin-3A, LPP-3A, and F-actin intensity at BCJs and MCJs pre- and post-extracellular ATP addition. Data from individual videos are shown as points with a LOESS regression function shown as a line. (*G)* Proposed model for the differential mechanoaccumulation of the different LIM proteins used in this study.

## Discussion

In this study, we use the embryonic epithelium in developing *Xenopus laevis* embryos to examine the mechanism by which LIM domain-containing proteins, which are known to stabilize and repair strained F-actin filaments, are recruited to TCJs as mechanical force increases. We found that both Zyxin and LPP are primarily recruited to TCJs through their LCRs, yet the N-terminus plays a regulatory role in the extent and kinetics of the mechanoaccumulation (**Fig. 6 *G***).

Our experimental data and docking simulations demonstrate that Zyxin and LPP respond to strained actin with distinct mechanisms. Our data suggests that Zyxin is a first responder at strained actin, as Zyxin mechanoaccumulation exhibits an early peak but then plateaus (**Fig. 2**). Following this initial recruitment, both LPP and Zyxin may bind and stabilize cracked actin. Surprisingly, the docking simulations suggest that Zyxin’s LCR has a weaker binding interaction with cracked actin (**Fig. 5**) which may lead to more transient interactions at the crack site. Our modeling focuses on the potential binding interactions between Zyxin or LPP’s LCR and the F-actin crack sites but cannot explain the differential mechanoaccumulation we observed: Zyxin mechanoaccumulates at TCJs to a greater extent than LPP. LPP’s higher binding strength for the crack sites suggests that the interactions between LCRs and the crack sites are not the only factor that drives the observed difference in mechanoaccumulation at TCJs (**Fig. 2**). A previous docking study of isolated LIM domains on cracked actin demonstrated interactions between LIM domains and other sites on the strained F-actin filament, including the ends of the filament (33). At TCJs, F-actin filament ends may be anchored to cell-cell junctions by scaffolding proteins, making TCJs a region of highly concentrated F-actin filament ends. It is plausible that binding interactions between Zyxin’s LCR and F-actin filament ends might underlie the observed difference between Zyxin and LPP’s mechanoaccumulation. Alternatively, the way acute mechanical force was applied to the *Xenopus* embryonic epithelium (hypoosmotic conditions + extracellular ATP) might not strain F-actin filaments in the same way that our computational modeling does. F-actin dynamically responds to tension by changing conformation through twisting and bending in addition to crack site formation (55,56). Future work is needed to predict the binding between Zyxin and LPP’s LCRs and actin filaments while accounting for multiple conformational changes to F-actin. Thus, Zyxin and LPP’s differential sensing of strained F-actin suggests that different LIM domain-containing proteins could serve as nuanced regulators of F-actin stability.

Our previous work showed that Zyxin and LPP preferentially localize to sites of stable F-actin (39). However, it remained unknown whether this was due to Zyxin and LPP stabilizing F-actin or to their recruitment to sites of stable F-actin. The current study shows that Zyxin and LPP are recruited to sites of high tension simultaneously with F-actin and maintain their localization as F-actin intensity plateaus. This pattern of mechanoaccumulation suggests that Zyxin and LPP sense and bind to strained F-actin, then stabilize it. This is consistent with the prevailing model for Zyxin and LPP recruitment and subsequent reinforcement of actin at stress fibers (29,31,34). Interestingly, Zyxin and LPP maintain their localization after F-actin intensity plateaus, suggesting that once the proteins are present at F-actin, they continue to stabilize the filaments even after the initial increase in force.

Our data points to a relationship between actomyosin-generated tension and mechanoaccumulation of Zyxin and LPP. Several studies have found a correlation between Zyxin family members and tissue tension, which suggests that the Zyxin family protein mechanoaccumulation on strained actin may both respond to increased tension and lead to better transmission of tension. For example, a recent study found that knocking down Zyxin decreased junctional tension at AJs in *Drosophila* larval wing imaginal discs (57). Additionally, another Zyxin family LIM domain-containing protein, Wtip, is recruited to cell-cell junctions in a tension-sensitive manner and is necessary for apical constriction of the *Xenopus* neural tube (58). Future studies should investigate how Zyxin and/or LPP’s mechanoaccumulation on and regulation of actomyosin associated with TCJs contributes to morphogenesis events involving high physiological forces such as neural tube closure.

## Methods

Videos were acquired with Airyscan or standard confocal microscopy, and image analysis was performed using FIJI and the FIJI Plugin, Trainable Weka Segmentation. Normalization and statistical analyses were performed using Microsoft Excel or RStudio. LCR and cracked F-actin docking models were generated using HADDOCK. Details of all methods and protocols are found in the *SI Appendix*.

## Acknowledgements

We thank the Liu lab, Miller lab, and Shiladitya Banerjee (Georgia Institute of Technology) for helpful discussions and feedback on this research. We acknowledge the University of Michigan BRCF Microscopy Core (RRID: SCR 026722) for training and use of the Zeiss LSM 980 with Airyscan2 confocal microscope. We are grateful to the NIH-funded National *Xenopus* Resource (RRID: SCR_013731) and Xenbase (RRID: SCR_003280) for public support of *Xenopus* research.

## AI Acknowledgement

During analysis of the data, the authors used the University of Michigan’s GPT and MATLAB Copilot Chat to draft and debug code to analyze, normalize, and plot the quantified data. After using the tool, the authors reviewed and edited the code as needed and take full responsibility for the publication’s content.

## Author Contributions

Conceptualization, K.K.T., A.L.M., A.P.L.;

Methodology, K.K.T., B.A., T.P., K.M.H., S.S.I., A.L.M., A.P.L.;

Formal Analysis, K.K.T., L.M.B., Y.B., T.P.;

Investigation, K.K.T., B.A., T.P.; Resources, K.K.T., L.X., A.L.M., A.P.L.;

Writing, K.K.T., B.A., T.P., Y.B.;

Writing – review and editing, K.K.T., T.P., B.A., Y.B., K.M.H., S.S.I., L.M.B., L.X., G.A.V., A.L.M., A.P.L.;

Visualization, K.K.T., B.A., T.P., Y.B.;

Supervision, K.K.T., G.A.V., A.L.M., A.P.L.;

Funding acquisition, K.K.T., G.A.V., A.L.M., A.P.L.

## References

1. Charras, G., & Yap, A. (2018). Tensile forces and mechanotransduction at cell–cell junctions. Current Biology, 28(8), R445–R457.

2. Perrin, L., & Vignjevic, D. M. (2023). The emerging roles of the cytoskeleton in intestinal epithelium homeostasis. Semin Cell Dev Biol, 150-151:23–27.

3. Schell, C., & Huber, T. B. (2017). The evolving complexity of the podocyte cytoskeleton. Journal of the American Society of Nephrology, 28(11), 3166–3174.

4. Trichas, G., Smith, A. M., White, N., Wilkins, V., Watanabe, T., Moore, A., … & Srinivas, S. (2012). Multi-cellular rosettes in the mouse visceral endoderm facilitate the ordered migration of anterior visceral endoderm cells. PLoS biology, 10(2), e1001256.

5. Higashi, T., & Miller, A. L. (2017). Tricellular junctions: how to build junctions at the TRICkiest points of epithelial cells. Molecular biology of the cell, 28(15), 2023–2034.

6. Higashi, T., & Chiba, H. (2020). Molecular organization, regulation and function of tricellular junctions. Biochimica et Biophysica Acta (BBA)-Biomembranes, 1862(2), 183143.

7. Bosveld, F., & Bellaïche, Y. (2020). Tricellular junctions. Current Biology, 30(6), R249–R251.

8. Arnold T.R., Stephenson R.E., Miller, A,L. (2017). Rho GTPases and actomyosin: partners in regulating epithelial cell-cell junction structure and function. Experimental cell research, 1;358(1):20–30.

9. Hartsock, A. and Turner J.R. (2008). Adherens and tight junctions: structure, function and connections to the actin cytoskeleton. Biochem Biophys Acta, 1778(3): p. 660–9.

10. Yonemura, S. (2011). Cadherin-actin interactions at adherens junctions. Curr. Opin. Cell Biol. 23,515–522.

11. Choi, W.et al. (2016). Remodeling the zonula adherens in response to tension and the role of afadin in this response. J. Cell Biol. 213,243–260.

12. Gomez, G. A., Mclachlan, R. W. & Yap, A. S. (2011). Productive tension: force-sensing and homeostasis of cell–cell junctions. Trends Cell Biol. 21,499–505.

13. Yonemura, S., Wada, Y., Watanabe, T., Nagafuchi, A., & Shibata, M. (2010). α-Catenin as a tension transducer that induces adherens junction development. Nature cell biology, 12(6), 533–542.

14. Marie, H., Pratt, S. J., Betson, M., Epple, H., Kittler, J. T., Meek, L., … & Braga, V. M. (2003). The LIM protein Ajuba is recruited to cadherin-dependent cell junctions through an association with α-catenin. Journal of Biological Chemistry, 278(2), 1220–1228.

15. Alégot, H., Markosian, C., Rauskolb, C., Yang, J., Kirichenko, E., Wang, Y. C., & Irvine, K. D. (2019). Recruitment of Jub by α-catenin promotes Yki activity and Drosophila wing growth. Journal of Cell Science, 132(5), jcs222018.

16. Yu, H. H., & Zallen, J. A. (2020). Abl and Canoe/Afadin mediate mechanotransduction at tricellular junctions. Science, 370(6520).

17. Masuda, S., Oda, Y., Sasaki, H., Ikenouchi, J., Higashi, T., Akashi, M., … Furuse, M. (2011). LSR defines cell corners for tricellular tight junction formation in epithelial cells. Journal of Cell Science, 124(4), 548–555.

18. Cho, Y., Haraguchi, D., Shigetomi, K., Matsuzawa, K., Uchida, S., & Ikenouchi, J. (2022). Tricellulin secures the epithelial barrier at tricellular junctions by interacting with actomyosin. Journal of Cell Biology, 221(4).

19. Campbell, H. K., Maiers, J. L., & DeMali, K. A. (2017). Interplay between tight junctions & adherens junctions. Experimental cell research, 358(1), 39–44.

20. Adhikary, B., Chang, A., Higashi, A. Y., Chiba, H., Miller, A. L., & Higashi, T. (2026). PAK4 promotes vertex remodeling to maintain epithelial integrity and barrier function. Journal of Cell Biology, 225(9), e202510024.

21. Spadaro, D., Le, S., Laroche, T., Mean, I., Jond, L., Yan, J., & Citi, S. (2017). Tension-dependent stretching activates ZO-1 to control the junctional localization of its interactors. Current biology, 27(24), 3783–3795.

22. 22. van den Goor, L., Iseler, J., Koning, K. M., & Miller, A. L. (2024). Mechanosensitive recruitment of Vinculin maintains junction integrity and barrier function at epithelial tricellular junctions. Current Biology, 34(20), 4677–4691.

23. Jacobs, T., Sanchez, J. I., Reger, S., & Luschnig, S. (2025). Rho/Rok-dependent regulation of actomyosin contractility at tricellular junctions restricts epithelial permeability in Drosophila. Current Biology, 35(6), 1181–1196.

24. Prudhomme, I. S., Brooks, E. R., Taneja, N., Bhattacharya, B., LaFleche, B. J., Furuta, Y., & Zallen, J. A. (2026). Genetically engineered ESC-derived embryos reveal Vinculin-dependent force responses required for mammalian neural tube closure. eLife15:RP110421.

25. Zheng, Q. H., Zhang, C., Wang, M. X., Xiang, X., Zhang, S., Wang, Y., & Yu, H. H. (2026). Edge-vertex flow enables rapid adhesion reinforcement under tension. Journal of Cell Biology, 225(6), e202601182.

26. Anderson, C. A., Kovar, D. R., Gardel, M. L. and Winkelman, J. D. (2021). LIM domain proteins in cell mechanobiology. Cytoskeleton, 78(6), pp.303–311.

27. Siddiqui, M., Badmalia, M., & Patel, T. (2021). Bioinformatic Analysis of Structure and Function of LIM Domains of Human Zyxin Family Proteins. International Journal of Molecular Sciences, 22(5), 2647.

28. Sala, S., & Oakes, P. W. (2023). LIM domain proteins. Current Biology, 33(9), R339–R341.

29. Sun, X., Phua, D. Y. Z., Axiotakis, L., Smith, M. A., Blankman, E., Gong, R., … Alushin, G. M. (2020). Mechanosensing through Direct Binding of Tensed F-Actin by LIM Domains. Developmental Cell, 55(4), 468–482.e7.

30. Winkelman, J. D., Anderson, C. A., Suarez, C., Kovar, D. R., & Gardel, M. L. (2020). Evolutionarily diverse LIM domain-containing proteins bind stressed actin filaments through a conserved mechanism. Proceedings of the National Academy of Sciences, 117(41), 25532–25542.

31. Call, G. S., Chung, J. Y., Davis, J. A., Price, B. D., Primavera, T. S., Thomson, N. C., … & Hansen, M. D. (2011). Zyxin phosphorylation at serine 142 modulates the zyxin head–tail interaction to alter cell–cell adhesion. Biochemical and biophysical research communications, 404(3), 780–784.

32. Phua, D. Y., Sun, X., & Alushin, G. M. (2024). Force-activated zyxin assemblies coordinate actin nucleation and crosslinking to orchestrate stress fiber repair. Current Biology, 35(4), 854–870.

33. Zsolnay, V., Gardel, M. L., Kovar, D. R., & Voth, G. A. (2024). Cracked actin filaments as mechanosensitive receptors. Biophysical Journal, 123(19), 3283–3294.

34. Hoffman, L. M., Jensen, C. C., Chaturvedi, A., Yoshigi, M., & Beckerle, M. C. (2012). Stretch-induced actin remodeling requires targeting of zyxin to stress fibers and recruitment of actin regulators. Molecular biology of the cell, 23(10), 1846–1859.

35. Katsuta, H., Okuda, S., Nagayama, K., Machiyama, H., Kidoaki, S., Kato, M., … Hirata, H. (2023). Actin crosslinking by α-actinin averts viscous dissipation of myosin force transmission in stress fibers. IScience, 26(3), 106090.

36. 36. Brühmann, S., Ushakov, D. S., Winterhoff, M., Dickinson, R. B., Curth, U., & Faix, J. (2017). Distinct VASP tetramers synergize in the progressive elongation of individual actin filaments from clustered arrays. PNAS, E5815-E5824.

37. Lynch, A. M., Zhu, Y., Lucas, B. G., Winkelman, J. D., Bai, K., Martin, S. C., … & Hardin, J. (2022). TES-1/Tes and ZYX-1/Zyxin protect junctional actin networks under tension during epidermal morphogenesis in the C. elegans embryo. Current Biology, 32(23), 5189–5199.

38. Slabodnick, M. M., Tintori, S. C., Prakash, M., Zhang, P., Higgins, C. D., Chen, A. H., … & Goldstein, B. (2023). Zyxin contributes to coupling between cell junctions and contractile actomyosin networks during apical constriction. PLoS genetics, 19(3), e1010319.

39. Tjoelker K.K., Bashirzadeh Y., Beel L.M., Liu A.P., Miller A.L. (2026). LIM domain-containing proteins Zyxin and LPP localize to apical epithelial cell-cell junctions at regions of stable F-actin. microPublication Biology. 10.17912/micropub.biology.002233.

40. Hansen, M. D., & Beckerle, M. C. (2006). Opposing roles of zyxin/LPP ACTA repeats and the LIM domain region in cell-cell adhesion. Journal of Biological Chemistry, 281(23), 16178–16188.

41. 41. von Dassow, M., & Davidson, L. A. (2009). Natural variation in embryo mechanics: gastrulation in Xenopus laevis is highly robust to variation in tissue stiffness. Developmental dynamics, 238(1), 2–18.

42. Pinheiro, D., Hannezo, E., Herszterg, S., Bosveld, F., Gaugue, I., Balakireva, M., … & Bellaïche, Y. (2017). Transmission of cytokinesis forces via E-cadherin dilution and actomyosin flows. Nature, 545(7652), 103–107.

43. 43. Landino, J., Misterovich, E., van den Goor, L., Adhikary, B., Chumki, S., Davidson, L. A., & Miller, A. L. (2025). Neighbor cells restrain furrowing during Xenopus epithelial cytokinesis. Developmental cell, 60(16), 2139–2148.

44. Luu, O., David, R., Ninomiya, H., & Winklbauer, R. (2011). Large-scale mechanical properties of Xenopus embryonic epithelium. Proceedings of the National Academy of Sciences, 108(10), 4000–4005.

45. Javed, A., Stubb, A., Villeneuve, C., Myllymäki, S. M., Peters, F., Rübsam, M., … & Wickström, S. A. (2025). Piezo1 balances membrane and cortex tension to stabilize intercellular junctions and maintain the epithelial barrier. Journal of Cell Science, 138(16), jcs263938.

46. Kim, Y., Hazar, M., Vijayraghavan, D.S., Song, J., Jackson, T.R., Joshi, S.D., Messner, W.C., Davidson, L.A., and LeDuc, P.R. (2014). Mechanochemical actuators of embryonic epithelial contractility. Proc Natl Acad Sci U S A 111, 14366–14371.

47. Arnold, T.R., Shawky, J.H., Stephenson, R.E., Dinshaw, K.M., Higashi, T., Huq, F., Davidson, L.A., and Miller, A.L. (2019). Anillin regulates epithelial cell mechanics by structuring the medial-apical actomyosin network. Elife 8:e39065.

48. 48. Joshi, S.D., von Dassow, M., and Davidson, L.A. (2010). Experimental control of excitable embryonic tissues: three stimuli induce rapid epithelial contraction. Exp Cell Res 316, 103–114.

49. Joshi, S. D., Jackson, T. R., Zhang, L., Stuckenholz, C., & Davidson, L. A. (2025). Supracellular contractility in Xenopus embryo epithelia regulated by extracellular ATP and the purinergic receptor P2Y2. Journal of cell science, 138(18), jcs263877.

50. Phillips, T. A., Marcotti, S., Cox, S., & Parsons, M. (2024). Imaging actin organisation and dynamics in 3D. Journal of Cell Science, 137(2), jcs261389.

51. Dominguez, C., Boelens, R., Bonvin, A. M. J. J. (2003). HADDOCK: a protein–protein docking approach based on biochemical or biophysical information. J. Am. Chem. Soc. 125, 1731–1737. (10.1021/ja026939x)

52. 52. van Zundert, G. C. P., Rodrigues, J. P. G. L. M., Trellet, M., Schmitz, C., Kastritis, P. L., Karaca, E., Melquiond, A. S. J., van Dijk, M., de Vries, S. J., Bonvin, A. M. J. J. (2016). The HADDOCK2.2 web server: user-friendly integrative modeling of biomolecular complexes. J. Mol. Biol. 428(4), 720–725. (10.1016/j.jmb.2015.09.014).

53. Weng, G., Wang, E., Wang, Z., Liu, H., Zhu, F., Li, D., Hou, T. (2019). HawkDock: a web server to predict and analyze the protein–protein complex based on computational docking and MM/GBSA. Nucleic Acids Res. 47(W1), W322–W330. (10.1093/nar/gkz397).

54. Massova, I., Kollman, P. A. (2000). Combined molecular mechanical and continuum solvent approach (MM-PBSA/GBSA) to predict ligand binding. Perspect. Drug Discov. Des. 18, 113–135. (10.1023/A:1008763014207)

55. Nakamura, M., Hui, J., & Parkhurst, S. M. (2023). Bending actin filaments: twists of fate. Faculty Reviews, 12, 7.

56. Banerjee, D. S., Freedman, S. L., Murrell, M. P., & Banerjee, S. (2024). Growth-induced collective bending and kinetic trapping of cytoskeletal filaments. Cytoskeleton, 81(8), 409–419.

57. Singh, H., Brooks, E., Jinnai, K., Kondo, S., Manning, S. A., Kroeger, B., & Harvey, K. F. (2026). The mechanosensitive protein Zyxin influences Hippo signaling and tissue growth via adherens junctions and basal spot junctions in Drosophila. Current Biology, 36, 4100–4112.e4.

58. Chu, C. W., Xiang, B., Ossipova, O., Ioannou, A., & Sokol, S. Y. (2018). The Ajuba family protein Wtip regulates actomyosin contractility during vertebrate neural tube closure. Journal of Cell Science, 131(10), jcs213884.

