## Supplemental Data and Methods for "Zyxin and LPP differ in their sensing of strained actin filaments at tricellular junctions"

**This PDF file includes:**

Figures S1 to S5
Table S1 to S2
Materials and Methods
Legends for Movies S1 to S10
SI References

### Figures

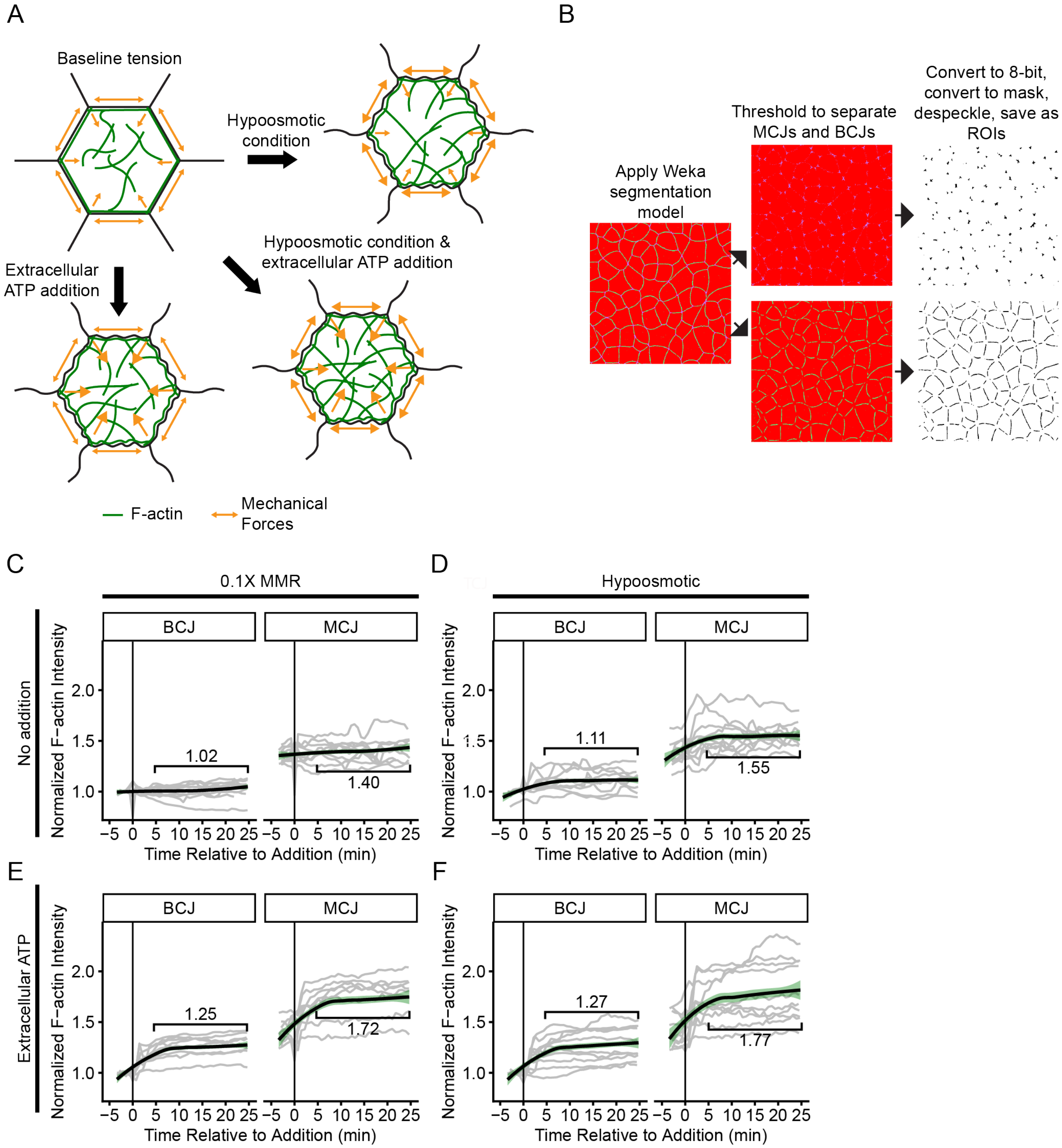


**Fig. S1. Extracellular ATP addition and hypoosmotic conditions result in an additive accumulation of F-actin at cell-cell junctions.**

A. Schematic of hypoosmotic condition and extracellular ATP additions. RNAse-free water and/or ATP is added to the *Xenopus* embryo during live confocal imaging.

B. Schematic of applying the Weka Segmentation model for segmenting images into ROIs for MCJs and BCJs. A Weka Segmentation model was trained on pre- and post-extracellular ATP addition images of ZO-1 from 12 videos. After applying the segmentation model, the MCJs and BCJs were separated using thresholding in FIJI. The thresholded images were then converted to 8-bit, converted to a mask, despeckled, and saved as ROIs.

C-F. Quantification of F-actin probe (LifeAct-miRFP703) intensity at BCJs and MCJs pre- and post-0.1X MMR addition (C), hypoosmotic condition (RNAse-free $H_2O$) (D), extracellular ATP in 0.1X MMR (E), or extracellular ATP in RNAse-free $H_2O$ (F). The measured signal was normalized to the average intensity at BCJs prior to addition. The averaged normalized intensity was calculated for 5 minutes to 25 minutes after the addition of the indicated condition. Regression line shows the LOESS fit.

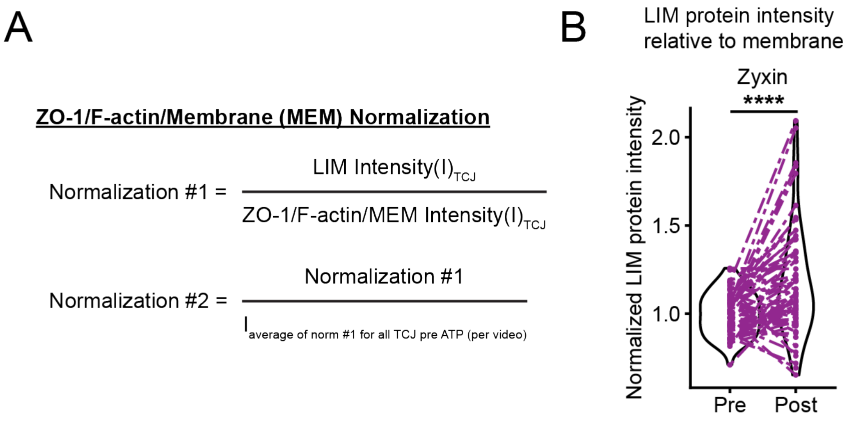


**Fig. S2. LIM protein intensity quantification and normalizations.**

A. Formulas for normalizing LIM protein intensity at TCJs relative to ZO-1, F-actin, or membrane intensity.

B. Quantification of membrane-normalized Zyxin-mNeonGreen intensity at TCJs pre- and post-extracellular ATP addition. Statistics: paired *t*-test; n = 7 clutches, 12 embryos, 84 TCJs; ****$p \leq 0.0001$.

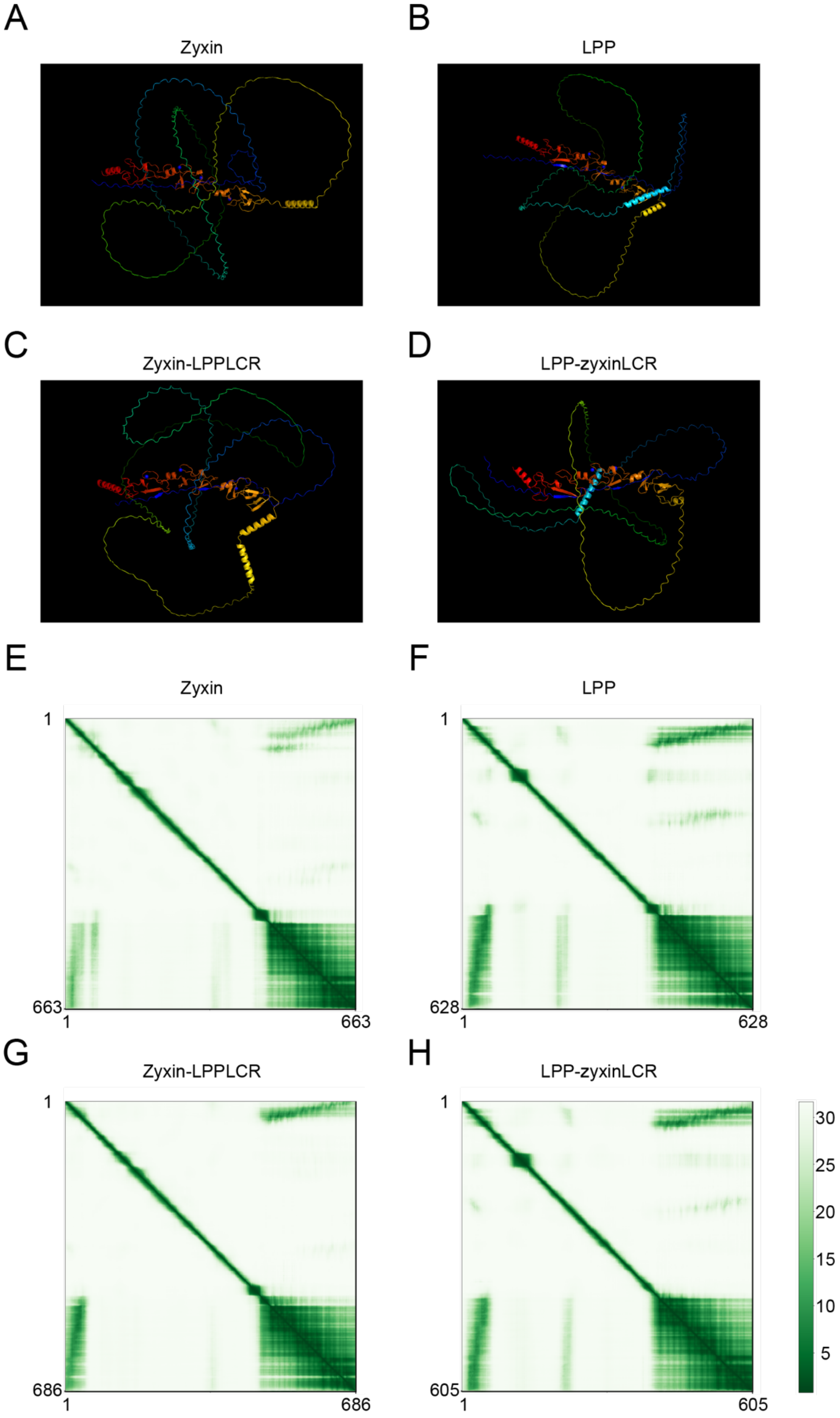


**Fig. S3. Predicted head-tail interactions for Zyxin and LPP: Zyxin has a less confident head-tail interaction prediction than LPP or chimeras.**

A-D. AlphaFold3 structural prediction of Zyxin (A), LPP (B), Zyxin-LPPLCR (C), LPP-ZyxinLCR (D) with three $Zn^{2+}$ ions (the LCR region has zinc finger motifs that chelate zinc ions (1), so $Zn^{2+}$ ions were included in the predicted structure).

E-H. Predicted alignment error (PAE) plot of AlphaFold3 structural prediction of Zyxin (E), LPP (F), Zyxin-LPPLCR (G), LPP-ZyxinLCR (H) with three $Zn^{2+}$ ions.

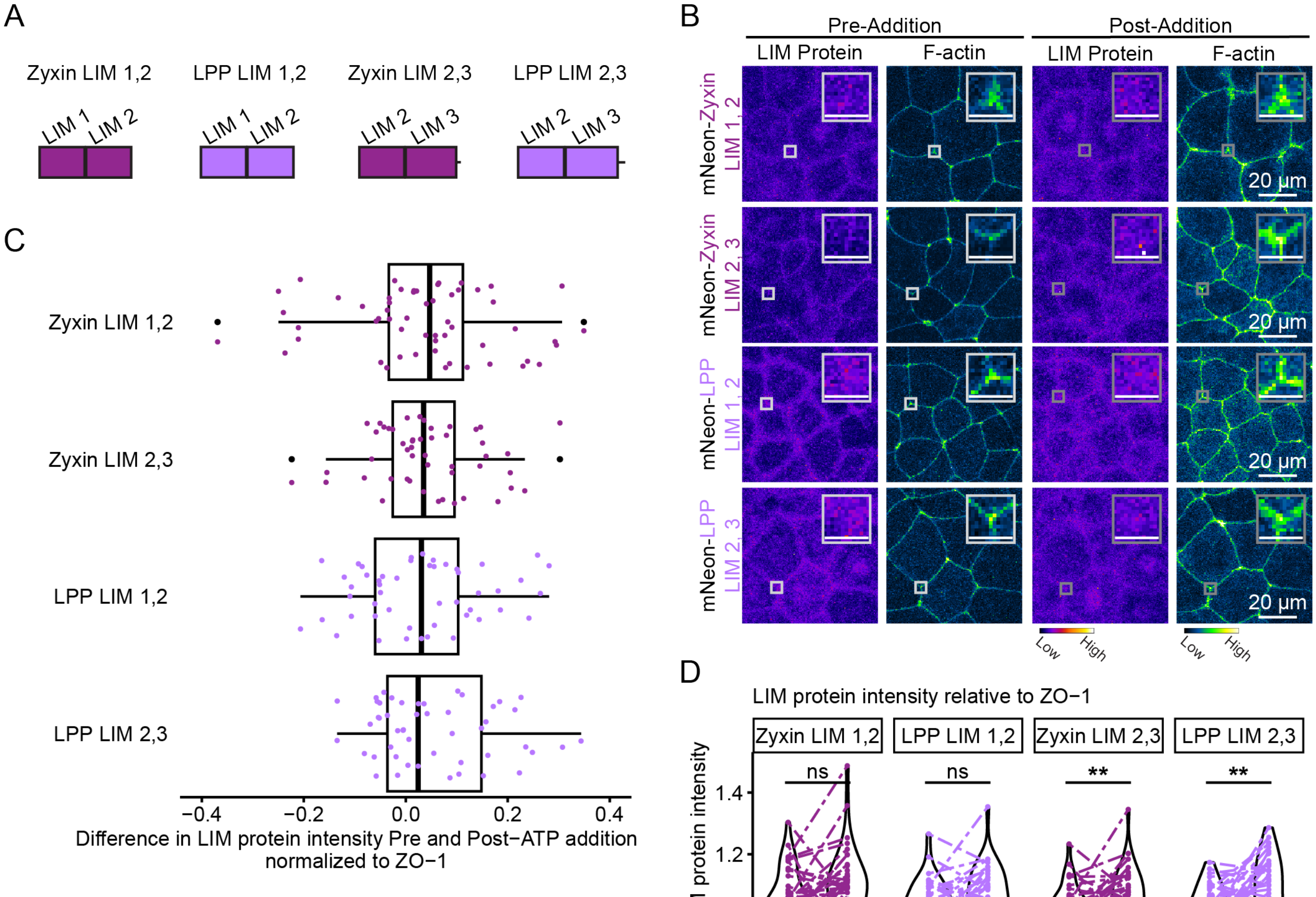


| | 1 | 2 | 3 | 4 |
|---|---|---|---|---|
| 1. Zyxin LIM 1,2 | X | ns | ns | ns |
| 2. Zyxin LIM 2,3 | ns | X | ns | ns |
| 3. Zyxin LCR | ns | ns | X | ns |
| 4. LPP LCR | ns | ns | ns | X |

**Fig. S4. LIM deletions from Zyxin and LPP's LCRs nearly eliminate mechanoaccumulation.**

A. Domain diagram of LIM deletions from the LCRs.

B. Live confocal images of cells expressing Zyxin LIM 1,2 (mNeonGreen-ZyxinLIM1,2, FIRE LUT), Zyxin LIM 2,3 (mNeonGreen-Zyxin-LIM2,3, FIRE LUT), LPP LIM 1,2 (mNeonGreen-

LPPLIM1,2, FIRE LUT), LPP LIM 2,3 (mNeonGreen-LPPLIM2,3, FIRE LUT), and F-actin probe (LifeAct-miRFP703, GreenFireBlue LUT) pre- and post-extracellular ATP addition. The enlarged insets highlight changes in Zyxin LIM 1,2, Zyxin LIM 2,3, LPP LIM 1,2, LPP LIM 2,3, and F-actin mechanoaccumulation at TCJs. Scale bar, 20 μm. Inset scale bar, 5 μm.

C. Quantification of the difference in LIM protein intensity (normalized to ZO-1 intensity) pre- and post-extracellular ATP addition. Statistics (shown in chart, bottom) by ANOVA.

D. Quantification of Zyxin LIM 1,2, Zyxin LIM 2,3, LPP LIM 1,2, LPP LIM 2,3 intensity (normalized to ZO-1 intensity) at TCJs pre- and post-extracellular ATP addition. For Zyxin LIM 1,2, n = 2 clutches, 8 embryos, 56 TCJs; Zyxin LIM 2,3, n = 2, 7, 49; LPP LIM 1,2, n = 2, 7, 49; LPP LIM 2,3, n = 2, 7, 49. Statistics: paired *t*-test; $^{**}p \leq 0.01$.

A

```
                  1 [        .         .         .         .  ↓      :         .         .         . 80
zyxinLCR_human      --CGRCHQPLARAQPAVRALGQLFHIACFTCHQCAQQLQGQQFYSLEGAPYCEGCYTDTLEKCNTCGEPITDRMLRATGK
zyxinLCR_canine     --CGRCHQPLARTQPAVRALGQLFHITCFTCHQCGQQLQGQQFYSLEGAPYCEGCYTDTLEKCNTCGQPITDRMLRATGK
zyxinLCR_mouse      --CGKCNQPLARAQPAVRALGQLFHITCFTCHQCQQQLQGQQFYSLEGAPYCEGCYTDTLEKCNTCGQPITDRMLRATGK
zyxinLCR_Xenopus    --CGFCGRGLSRTETVVRAGEHLYHVACFTCSRCDQQLQGQQYYESAGKPLCDECYQDTLECCAVCDKKITERLLKAIGK
LPPLCR_human        GRCARCGENVVGEGTGCTAMDQVFHVDCFTCIICNNKLRGQPFYAVEKKAYCEPCYINTLEQCNVCSKPIMERILRATGK
LPPLCR_canine       GRCARCGENVVGEGTGCTAMDQVFHVDCFTCIICNNKLRGQPFYAVEKKAYCEPCYINTLEQCSVCSKPIMERILRATGK
LPPLCR_mouse        GRCARCGENVVGEGTGCTAMDQVFHVDCFTCIVCDVKLRGQPFYAVEKKAYCEPCYINTLEQCSVCSKPIMERILRATGK
LPPLCR_Xenopus      GRCSRCGENVVGEGTGCTAMDQVFHVECFTCMTCNSKLRGQPFYAVEKKAFCEPCYIRTLEKCNVCAKPIMERILRATGK
consensus/100%      ..CuhCtpslstttssspAhtplaHltCFTC..Ct.pLpGQ.aYt.tttshC-.CY.pTLEpCssCsp.Ih-RhL+AhGK
consensus/90%       ..CuhCtpslstttssspAhtplaHltCFTC..Ct.pLpGQ.aYt.tttshC-.CY.pTLEpCssCsp.Ih-RhL+AhGK
consensus/80%       ..Cu+CtpslstttsuspAhsQlFHlsCFTCh.CsppLpGQ.FYulEttsaCEsCYhsTLEpCssCupPIh-RhLRATGK
consensus/70%       ..CuRCspslstptsuspAhsQlFHlsCFTCh.CsppLpGQ.FYulEttsYCEsCYhsTLEpCssCucPIh-RhLRATGK

                 81          .       ↓ .         .         .         .         .         :         . 160
zyxinLCR_human      AYHPHCFTCVVCARPLEGTSFIVDQANRPHCVPDYHKQYAPRCSVCSEPIMPEPGRDETVRVVALDKNFHMKCYKCEDCG
zyxinLCR_canine     AYHPQCFTCVVCACPLEGTSFIVDQANRPHCVPDYHKQYAPRCSVCEGPIMPEPGREETVRVVALDKNFHMKCYKCEDCG
zyxinLCR_mouse      AYHPQCFTCVVCACPLEGTSFIVDQANQPHCVPDYHKQYAPRCSVCSEPIMPEPGRDETVRVVALDKNFHMKCYKCEDCG
zyxinLCR_Xenopus    SYHPSCFTCAVCKCSLQGEPFIVDDNKLPHCVNDYHRRYAPRCCVCGDPIAPEPGRDETVRVVALEKNFHMMCYKCEDCG
LPPLCR_human        AYHPHCFTCVMCHRSLDGIPFTVDAGGLIHCIEDFHKKFAPRCSVCKEPIMPAPGQEETVRIVALDRDFHVHCYRCEDCG
LPPLCR_canine       AYHPHCFTCVMCHRSLDGIPFTVDAGGLIHCIEDFHKKFAPRCSVCKEPIMPAPGQEETVRIVALDRDFHVHCYRCEDCG
LPPLCR_mouse        AYHPHCFTCVMCHRSLDGIPFTVDACGLIHCIEDFHKKFAPRCSVCKEPIMPAPGQEETVRIVALDRDFHVHCYRCEDCG
LPPLCR_Xenopus      AYHPHCFTCVVCFRSLDGVPFTVDASGQIHCIEDFHKKFAPRCSVCKEPIMPAPGQEETVRIVALDRDFHVQCYRCEDCG
consensus/100%      uYHPpCFTCshChpsLpG.sFhVDtst..HCl.DaH+paAPRCsVCttPIhPtPGp-ETVRlVAL-+sFHh.CY+CEDCG
consensus/90%       uYHPpCFTCshChpsLpG.sFhVDtst..HCl.DaH+paAPRCsVCttPIhPtPGp-ETVRlVAL-+sFHh.CY+CEDCG
consensus/80%       AYHPpCFTCVhCtpsL-GhsFhVDtss..HCl.DaHKpaAPRCSVCp-PIMPtPGp-ETVRlVALD+sFHhpCY+CEDCG
consensus/70%       AYHPpCFTCVhCtpsL-GhsFhVDtush.HCl.DaHKpaAPRCSVCpEPIMPtPGp-ETVRlVALD+sFHh+CY+CEDCG

                161          .  ↓      .         .         ] 200
zyxinLCR_human      KPLSIEADDNGCFPLDGHVLCRKCHTARAQT---------
zyxinLCR_canine     KPLSIEADDNGCFPLDGHVLCRKCHTARAQT---------
zyxinLCR_mouse      KPLSIEADDNGCFPLDGHVLCRKCHSARAQT---------
zyxinLCR_Xenopus    CPLSIEADDAGCFPLDGHVLCKKCHTVRARAALG------
LPPLCR_human        GLLS-EGDNQGCYPLDGHILCKTCNSARIRVLTAKASTDL
LPPLCR_canine       GLLS-EGDNQGCYPLDGHILCKTCNSARIRVLTAKASTDL
LPPLCR_mouse        GLLS-EGDNQGCYPLDGHILCKTCNSARIRVLTAKASTDL
LPPLCR_Xenopus      SLLS-EGENQGCYPLDGHILCKACNAVRIQALTAKSSTDL
consensus/100%      t.LS.Eu-stGCaPLDGHlLC+tCpssRhps.........
consensus/90%       t.LS.Eu-stGCaPLDGHlLC+tCpssRhps.........
consensus/80%       t.LS.EuDspGCaPLDGHlLC+pCposRhps.........
consensus/70%       t.LS.EuDspGCaPLDGHlLC+pCpoARhps.........
```

B

zyxin

| Similarity human/*Xenopus* |
|---|
| 49.3% |

| Domain | Similarity human/*Xenopus* |
|---|---|
| α-actinin binding | 61.6% |
| ActA repeats | 53.8% |
| LCR | 75.5% |

LPP

| Similarity human/*Xenopus* |
|---|
| 74.3% |

| Domain | Similarity human/*Xenopus* |
|---|---|
| α-actinin binding | 86.4% |
| ActA repeats | 58.7% |
| LCR | 95.0% |

C

| Protein | Aromatic Residue | LIM domain in active pose | Crack Site | Residue Contribution |
|---|---|---|---|---|
| Zyxin | Y512 | LIM1 | CS2 | -3.30 kcal/mol |
| Zyxin | F642 | LIM3 | CS2 | -2.40 kcal/mol |
| LPP | Y601 | LIM3 | CS1 | -1.89 kcal/mol |

**Fig. S5. Zyxin and LPP sequence comparison highlighting conserved aromatic residues.**

A. Alignment of the sequences of human, canine, mouse, and *Xenopus* Zyxin and LPP LCRs (2). Conserved phenylalanine (or tyrosine) residues are highlighted in orange.

B. Left: Percent similarity of full-length Zyxin and its alpha-actinin binding site, ActA repeats, and LCR comparing human and *Xenopus* sequences. Right: Percent similarity of full-length LPP and its alpha-actinin binding site, ActA repeats, and LCR comparing human and *Xenopus* sequences.

C. Contributions for Zyxin Y512, Zyxin F642, and LPP Y601 in simulated docking between the LCR and cracked F-actin.

### Tables

**Table S1.** Cross-domain contributions to the binding energy between the LCRs and the F-actin crack sites.

| Protein | Crack Site | LIM domain in active pose | Residue | Residue's LIM domain | Actin monomer contact |
|---|---|---|---|---|---|
| LPP | CS1 | LIM1 | R487 | LIM2 | E361 and E364 actin monomer 'i-2' |
| LPP | CS2 | LIM1 | R487 | LIM2 | E117 actin monomer 'i-1' |
| LPP | CS2 | LIM1 | K497 | LIM2 | D80 and E83 actin monomer 'i-1' |
| LPP | CS2 | LIM1 | K491 | LIM2 | E125 actin monomer 'i-1' |
| LPP | CS2 | LIM1 | K546 | LIM2 | E2 actin monomer 'i-3' |
| LPP | CS1 & CS2 | LIM3 | R551 | LIM2 | Longer-range electrostatics 9-12 Å from actin |
| Zyxin | CS1 | LIM1 | F570 | LIM2 | E99 actin monomer 'i' |
| Zyxin | CS2 | LIM3 | R591 | LIM2 | S239 actin monomer 'i' |

**Table S2.** Constructs used to transcribe mRNA, cloning methods and primers used for generating constructs, and amount of mRNA per injection.

| Construct | Cloning Method | Forward Primer | Reverse Primer | mRNA/injection (pg) |
|---|---|---|---|---|
| pCS2+/Zyxin-LPPLCR-mNeonGreen | G block + Gibson Reaction | NA | NA | 10 pg |
| pCS2+/LPP-ZyxinLCR-mNeonGreen | G block + Gibson Reaction | NA | NA | 10 pg |
| pCS2+/mNeonGreen-ZyxinLCR | HiFi (NEBuilder) | gacgagctgtacaagagatcctgtggcttctgtgggcg | ctaatagttctagactgcagttatcccagggcagcgcg | 10 pg |
| pCS2+/mNeonGreen-LPPLCR | HiFi (NEBuilder) | gacgagctgtacaagagatctggacgctgctctcgctg | ctaatagttctagactgcagttacaagtcggtgcttgacttgg | 10 pg |
| pCS2+/ZyxinNterm-mNeonGreen | HiFi (NEBuilder) | gctacttgttctttttgcagaccatggacccagcggct | gagaggccttgaattcgaatttgagttccatagagtgcgcctc | 10 pg |

| | | | | |
|---|---|---|---|---|
| pCS2+/LPPNterm-mNeonGreen | HiFi (NEBuilder) | gctacttgttctttttgcagaccatgtctcatccatcatg | gagaggccttgaattcgaatccaaagtattcctcagaagg | 10 pg |
| pCS2+/Zyxin-3A-mNeonGreen | Mutagenesis (NEBaseChanger) | ggggcagcaggcctacgagagcg | tggagctgctgatcacac | 10 pg |
| pCS2+/LPP-3A-mNeonGreen | G block + restriction enzyme cloning | NA | NA | 10 pg |
| pCS2+/mNeonGreen-ZyxinLIM1,2 | G block + HiFi (NEBuilder) | NA | NA | 10 pg |
| pCS2+/mNeonGreen-LPPLIM1,2 | G block + HiFi (NEBuilder) | NA | NA | 10 pg |
| pCS2+/mNeonGreen-ZyxinLIM2,3 | G block + HiFi (NEBuilder) | NA | NA | 10 pg |
| pCS2+/mNeonGreen-LPPLIM2,3 | G block + HiFi (NEBuilder) | NA | NA | 10 pg |

### Supplementary Materials and Methods

#### *Xenopus laevis* model system

All experiments were carried out in compliance with the University of Michigan Institutional Animal Care and Use Committee and the U.S. Department of Health and Human Services Guide for the Care and Use of Laboratory Animals. Female *Xenopus laevis* frogs (*Xenopus* 1 or National *Xenopus* Resource [NXR]) were injected with human chorionic gonadotrophin to induce them to lay eggs. The eggs were collected, fertilized *in vitro*, dejellied in 2% cystine, pH 7.8 in 1X Mark's Modified Ringer's solution (MMR) (5 mM HEPES, 100 mM NaCl, 2 mM KCl, 1 mM $MgCl_2$, 2 mM $CaCl_2$, pH 7.4). Following dejellying, the embryos were transferred to 0.1X MMR.

At 2-cell or 4-cell stage, the embryos were microinjected with mRNA to express fluorescently-tagged proteins of interest. 5 nl was injected into the animal hemisphere of the embryos four times. Each 5 nl injection contained the following amount of mRNA: 10 pg or 20 pg pCS2+/Zyxin.S-mNeonGreen; 10 pg or 20 pg pCS2+/LPP.L-mNeonGreen; 10 pg pCS2+/Zyxin-LPPLCR-mNeonGreen; 10 pg pCS2+/LPP-ZyxinLCR-mNeonGreen; 10 pg mNeonGreen-ZyxinLCR; 10 pg mNeonGreen-LPPLCR; 10 pg pCS2+/ZyxinNterm-mNeonGreen; 10 pg pCS2+/LPPNterm-mNeonGreen; 10 pg pCS2+/Zyxin-3A-mNeonGreen; 10 pg pCS2+/LPP-3A-mNeonGreen; 10 pg pCS2+/mNeonGreen-ZyxinLIM1,2; 10 pg pCS2+/mNeonGreen-ZyxinLIM2,3; 10 pg pCS2+/mNeonGreen-LPPLIM1,2; 10 pg pCS2+/mNeonGreen-LPPLIM2,3; 150 pg pCSf107mT/LifeAct-miRFP703; 125 pg pCS2+/mRFP-ZO-1; 70 pg pCS2+/BFP-ZO-1. Embryos developed overnight at 15ºC until they reached blastula (Nieuwkoop and Faber stages 9-9.5) or gastrula stage (Nieuwkoop and Faber stages 10-11).

#### DNA constructs

New DNA constructs were cloned using primers described in *Table 2*. The primers were generated using NEBuilder Assembly Tool (https://nebuilder.neb.com/#!/) or NEBaseChanger (https://nebasechanger.neb.com/), as indicated in the table. G blocks were synthesized by Twist Bioscience. Constructs were verified by sequencing (Plasmidsaurus, Eugene OR, and Eurofin, Lancaster, PA).

All other DNA constructs were previously described: pCS2+/*Xenopus laevis* Zyxin.S-mNeonGreen; Tjoelker et al. 2026 (3); pCS2+/Xenopus laevis LPP.L-mNeonGreen; Tjoelker et al. 2026 (3); pCS2+/mRFP-ZO-1; Higashi et al. 2016 (4); pCS2+/TagBFP-ZO-1; Stephenson et al. 2019 (5); pCSf107mT/Lifeact-miRFP703; Yamamoto et al. 2021 (6).

#### mRNA preparation

mRNA was transcribed *in vitro* by first linearizing pCS2+ DNA constructs using Not1-HF. pCSf107mT/LifeAct-miRFP703 was not linearized. The DNA constructs were then transcribed with the mMessage mMachine SP6 Transcription Kit (Invitrogen) and purified with the RNeasy Mini Kit (Qiagen). mRNA was stored at -80ºC until use.

#### Image acquisition

Super-resolution live confocal imaging of blastula- or gastrula-stage *Xenopus* embryos in Figure 1 was performed using a Zeiss LSM 980 confocal microscope equipped with an Airyscan 2 detector and a Plan-Apochromat 63×/1.4 NA oil-immersion objective. Images were acquired with a 2.0X optical zoom. Z-stacks spanning approximately 8 µm were collected with a z-step size of 0.16 µm. Airyscan images were processed using Zeiss ZEN Blue software. The Wiener filter used for Airyscan processing was manually adjusted for each channel to optimize the signal-to-noise ratio. Blastula-stage embryos had been kept for 13-14 hours at 15°C and had not yet initiated dorsal blastopore closure. Mid- to late-gastrula-stage embryos were maintained for more than 15 hours at 15°C and were characterized by greater than 40% blastopore closure.

Images in Figures 2-4 and 6 and supplemental movies were captured using an inverted Olympus FV1000 confocal microscope with mFV-10-ASW software. Images were obtained with a supercorrected Plan Apo N 60XOSC objective (NA = 1.4, working distance = 0.12 mm). Embryos were mounted in a custom metal slide with a hole in the center; embryos were held in place with two coverslips attached to the metal slide with vacuum grease.

Time-lapse movies were acquired for mNeonGreen-tagged LIM protein, ZO-1 (BFP-ZO-1 or mRFP-ZO-1), and F-actin (LifeAct-miRFP703). The apical Z-planes were sequentially scanned using a 512 x 512-pixel area.

**Hypoosmotic condition and extracellular ATP addition**

Embryos were mounted in the custom metal slide by sandwiching them between two coverslips, but a portion of the chamber was left open with the top coverslip. After imaging 4-10 frames (pre-addition), 100 µl of one of the following was added in the opening: 0.1X MMR, RNAse-free $H_2O$, 500 µM ATP in 0.1X MMR, or 500 µM ATP in RNAse-free $H_2O$ and live imaging continued (post-addition).

**Quantification and Statistical Analysis**

Quantification was performed in Fiji on max-projected images, normalization in Microsoft Excel or RStudio, and statistical analysis and graphing in RStudio. Images in the figures were max-projected and manually adjusted to show relevant features in Fiji, and LUTs were applied as indicated in the figure legends.

**Quantification of Zyxin and LPP puncta**

For quantification of Zyxin and LPP puncta, TCJs were cropped, and Zyxin and LPP puncta at TCJs were counted manually in a blinded manner.

For quantification of super-resolution images, TCJs were cropped from a single z-plane. A single optical section was used to avoid artifacts from embryo drift during super-resolution imaging. The z-plane used for analysis was selected as the first apical slice showing uniform F-actin signal, typically approximately the fifth slice from the most apical plane, corresponding to a depth of approximately 0.64 µm given the 0.16 µm z-step size. For quantification of conventional confocal images in Figures 2 and 3, TCJs were cropped from a maximum-intensity projection.

**Quantification of TCJs before and after extracellular ATP addition**

For each embryo, images were max-projected. LIM protein, ZO-1, and F-actin intensities were measured in 5 µm circular regions of interest (ROIs) at seven TCJs and 3 cytosolic regions for pre- and post-extracellular ATP addition. One pre-ATP and one post-ATP frame were selected for in-focus signal and maximum cell contraction. LIM protein TCJ measurements were normalized to ZO-1, F-actin, or membrane signal in the same ROI. A paired t-test was performed to compare normalized pre- and post-ATP addition LIM protein intensity.

**Quantification of linescans that start along BCJs and pass through TCJs**

In Fiji, a line with a width of 0.828 µm was drawn along a BCJ, through a TCJ, then into the cytosol; the line was centered on the TCJ. The intensity values were normalized by the average of the lowest 20% of values in the linescan (cytosolic intensity).

**Quantification of BCJs and MCJs**

In Fiji, a Trainable Weka Segmentation model was trained using the ZO-1 channel for pre-ATP and post-ATP frames from 12 videos with the most extreme variation in ZO-1 intensity from the Zyxin and LPP datasets. The model was trained to identify BCJs and MCJs. The model was then applied to ZO-1 channels for videos with ATP. The model resulted in segmented time series. For the quantification of the dataset in *SI Appendix*, Fig. S1, which lacked the ZO-1 channel, a Trainable Weka Segmentation model was trained to identify BCJs and MCJs using 9 frames (3 time points from 3 videos) of the F-actin channel from the extracellular ATP in RNAse-free $H_2O$ data. The model was then applied to F-actin channel for the dataset and resulted in segmented time series.

Using the Threshold function, the segmentation for BCJs was separated from that of MCJs. For each BCJ and MCJ timeseries, the timeseries were turned into 8-bit, converted to a mask, and despeckled. The masks were converted into ROIs for measuring intensities in the LIM protein and F-actin channels for each time point.

In R, the normalized MCJ LIM protein and F-actin intensity values were averaged. For Figures 2, 4, and 6 the indicated values are for the averages for 5 to 10 minutes and 10 to 25 minutes following extracellular ATP addition. For *SI Appendix*, Fig. S1, the indicated values are for the average intensities from 5 to 25 minutes following extracellular ATP addition.

For the analysis of the kinetics, peaks in the time course plots were counted manually in a blinded manner. A contingency table was generated, and the odds ratio was calculated using the epitools R package.

**Batch particle image velocimetry**

We developed a custom algorithm that called the source code functions of the PIVlab (7) to measure the displacement field, divergence, and shear rate of fluorescent tags per pair of image frames using 4 passes of multi-pass fast-Fourier transform window deformation at interrogation window sizes of 64, 32, 16, and 8 in order. Overlap between interrogation windows was set to 50% for each pass. Velocity vectors were then validated with a standard deviation filter threshold of 8 and a local median filter threshold of 3.

Cell boundaries were first segmented using two-step Canny edge detection of contrast-adjusted protein intensity images, followed by custom image processing operations to eliminate morphological defects. To measure BCJ and MCJ dynamics, we developed custom Matlab functions that determine junction type by analyzing the labeled matrix of segmented cell boundary images within the tissue. To better distinguish between BCJ and MCJ behaviors, we assumed an 8 pixel-long PIV window belongs to MCJs when its shortest distance from at least 3 neighboring cells in all 8 directions is equal to or less than 3 windows long. A cell junction pixel was assumed to be BCJ when the pixel was, at maximum, 3 pixels away from a cell pair in all directions and further away from any other nearby cells.

Velocity magnitude, divergence, and shear rate of BCJs and MCJs were averaged per pair of images to obtain junction dynamics for each region over time. Since time points and intervals slightly differed across regions, we interpolated dynamic properties at fixed, rounded time points, obtained the means, and performed LOESS smoothing over the means per time point to obtain the total average of dynamic properties at each rounded time point (**Fig. 2 *F* and *G***). We measured the standard error from the mean at individual time points to demonstrate dynamic deviations across tissue regions.

**Zyxin/LPP LCR-cracked actin docking**

The structure of the cracked actin filament for all our docking calculations was obtained from the simulation of an actin filament under 300 pN tension (8). The *Xenopus laevis* Zyxin and LPP amino-acid sequences were obtained from UniProtKB entries A5H447 and Q52L12, respectively, and were used to generate the LCR structural models.

Zyxin or LPP LCRs were docked against the cracked actin models using HADDOCK (9-10), an information-driven protein-protein docking software that enables the use of user-defined interface restraints during complex generation. The two docking locations, crack site 1 (CS1) and crack site 2 (CS2), were defined using the same crack site residues defined in Zsolnay *et al*. (8). CS1 corresponds to the exposed interface between subdomain 2 (SD2) of monomer 'i' and SD1 and SD3 of monomer 'i-2', and CS2 corresponds to the secondary crack site interface located between SD4 of monomer 'i' and SD3 of monomer 'i-2' using the same monomer labeling scheme as our previous work.

For each Zyxin/LPP LCR and crack site combination, docking was performed separately by designating LIM1, LIM2, or LIM3 residues as binding residues, yielding 12 combinations in total (2 proteins x 3 LIM domains/protein x 2 crack sites/LIM domain). This arrangement helped us to assess how different LIM domains within the tandem LCR engage the cracked actin interface. Actin was treated as the receptor, and the Zyxin or LPP LCR as the ligand. For each HADDOCK run, docking clusters were ranked by the HADDOCK score (9-10), and the representative top-ranked pose from the best-scoring cluster was selected for post-docking energetic analysis. HADDOCK scores (9-10) each model using a weighted combination of van der Waals, electrostatic, desolvation, and restraint-violation terms (i.e., HADDOCK score = $E_{vdW} + 0.2E_{elec} + E_{desolv} + 0.1E_{AIR}$). $E_{AIR}$ is the penalty for violating the ambiguous interaction restraints derived from the specified active (and passive) interface residues. Active residues are expected to contact the active (or passive) surface patch on the partner, so models that fail to satisfy these restraints receive a larger positive $E_{AIR}$ value. This term is weighted by 0.1, allowing the restraints to guide docking without overwhelming the physical interaction terms. $E_{AIR}$ is therefore a restraint-consistency measure, not a physical binding energy.

Models with similar interface contacts are grouped into clusters, which are ranked using the average score of their four best members, and more negative scores indicate more plausible structures, while the Z-score indicates how strongly a cluster stands out from the others. The best representative static structure of the top-ranked cluster was carried forward for MM-GBSA calculations with HawkDock (11-12). It offers a web server for MM-GBSA-based binding-energy estimation and residue-wise energy decomposition for static protein-protein structures. MM-GBSA estimates relative binding free energy by combining molecular-mechanics interaction terms with implicit-solvent contributions. The reported MM-GBSA binding energies do not include the binding entropy term (-TΔS). Therefore, the reported total, calculated from vdW, elec, GB, and SA components, was interpreted as a relative energetic estimate rather than a complete absolute binding free energy. From HawkDock reported total energies, the nonpolar contribution was defined as vdW + SA. vdW reflects packing and dispersion interactions at the interface, whereas SA approximates the energetic contribution from burial of solvent-accessible nonpolar surface. The polar contribution was defined as elec + GB. Elec captures direct Coulombic attraction or repulsion, while GB represents changes in polar solvation and can offset favorable electrostatic contacts through desolvation penalties. The total energy was calculated as vdW + elec + GB + SA, with more negative values indicating more favorable relative binding; residue-wise decomposition was also used to identify energetic hotspots.

**Legends for Movies S1 to S10**

**Movie S1 (separate file). Zyxin and F-actin mechanoaccumulate at cell-cell junctions.** Zyxin-mNeonGreen (FIRE LUT) and Lifeact-miRFP703 (GreenFireBlue LUT) mechanoaccumulate at cell-cell junctions following addition of extracellular ATP (ATP addition is indicated by the magenta box). Scale bar, 20 µm. Playback at 5 fps.

**Movie S2 (separate file). LPP and F-actin mechanoaccumulate at cell-cell junctions.** LPP-mNeonGreen (FIRE LUT) and Lifeact-miRFP703 (GreenFireBlue LUT) mechanoaccumulate at cell-cell junctions following addition of extracellular ATP (ATP addition is indicated by the magenta box). Scale bar, 20 µm. Playback at 5 fps.

**Movie S3 (separate file). Zyxin-LPPLCR and F-actin mechanoaccumulate at cell-cell junctions.** Zyxin-LPPLCR-mNeonGreen (FIRE LUT) and Lifeact-miRFP703 (GreenFireBlue LUT) mechanoaccumulate at cell-cell junctions following addition of extracellular ATP (ATP addition is indicated by the magenta box). Scale bar, 20 µm. Playback at 5 fps.

**Movie S4 (separate file). LPP-ZyxinLCR and F-actin mechanoaccumulate at cell-cell junctions.** LPP-ZyxinLCR-mNeonGreen (FIRE LUT) and Lifeact-miRFP703 (GreenFireBlue LUT) mechanoaccumulate at cell-cell junctions following addition of extracellular ATP (ATP addition is indicated by the magenta box). Scale bar, 20 µm. Playback at 5 fps.

**Movie S5 (separate file). Zyxin LCR and F-actin mechanoaccumulate at cell-cell junctions.** ZyxinLCR-mNeonGreen (FIRE LUT) and Lifeact-miRFP703 (GreenFireBlue LUT) mechanoaccumulate at cell-cell junctions following addition of extracellular ATP (ATP addition is indicated by the magenta box). Scale bar, 20 µm. Playback at 5 fps.

**Movie S6 (separate file). LPP LCR and F-actin mechanoaccumulate at cell-cell junctions.** LPPLCR-mNeonGreen (FIRE LUT) and Lifeact-miRFP703 (GreenFireBlue LUT) mechanoaccumulate at cell-cell junctions following addition of extracellular ATP (ATP addition is indicated by the magenta box). Scale bar, 20 µm. Playback at 5 fps.

**Movie S7 (separate file). Zyxin's N-terminus and F-actin mechanoaccumulate at cell-cell junctions.** ZyxinNterm-mNeonGreen (FIRE LUT) and Lifeact-miRFP703 (GreenFireBlue LUT) mechanoaccumulate at cell-cell junctions following addition of extracellular ATP (ATP addition is indicated by the magenta box). Scale bar, 20 µm. Playback at 5 fps.

**Movie S8 (separate file). LPP's N-terminus and F-actin mechanoaccumulate at cell-cell junctions.** LPPNterm-mNeonGreen (FIRE LUT) and Lifeact-miRFP703 (GreenFireBlue LUT) mechanoaccumulate at cell-cell junctions following addition of extracellular ATP (ATP addition is indicated by the magenta box). Scale bar, 20 µm. Playback at 5 fps.

**Movie S9 (separate file). Zyxin-3A and F-actin mechanoaccumulate at cell-cell junctions.** Zyxin-3A-mNeonGreen (FIRE LUT) and Lifeact-miRFP703 (GreenFireBlue LUT) mechanoaccumulate at cell-cell junctions following addition of extracellular ATP (ATP addition is indicated by the magenta box). Scale bar, 20 µm. Playback at 5 fps.

**Movie S10 (separate file). LPP-3A and F-actin mechanoaccumulate at cell-cell junctions.** LPP-3A-mNeonGreen (FIRE LUT) and Lifeact-miRFP703 (GreenFireBlue LUT) mechanoaccumulate at cell-cell junctions following addition of extracellular ATP (ATP addition is indicated by the magenta box). Scale bar, 20 µm. Playback at 5 fps.

**SI References**


1. Anderson, C.A., Kovar, D.R., Gardel, M.L. and Winkelman, J.D. (2021). LIM domain proteins in cell mechanobiology. Cytoskeleton, 78(6), pp.303-311.
2. Madeira, F., Madhusoodanan, N., Lee, J., Eusebi, A., Niewielska, A., Tivey, A. R., ... & Butcher, S. (2024). The EMBL-EBI Job Dispatcher sequence analysis tools framework in 2024. Nucleic acids research, 52(W1), W521-W525.
3. Tjoelker K.K., Bashirzadeh Y., Beel L.M., Liu A.P., Miller A.L. (2026). LIM domain-containing proteins Zyxin and LPP localize to apical epithelial cell-cell junctions at regions of stable F-actin. microPublication Biology. 10.17912/micropub.biology.002233.
4. Higashi, T., Arnold, T. R., Stephenson, R. E., Dinshaw, K. M., & Miller, A. L. (2016). Maintenance of the epithelial barrier and remodeling of cell-cell junctions during cytokinesis. Current biology, 26(14), 1829-1842. (27345163)
5. Stephenson, R. E., Higashi, T., Erofeev, I. S., Arnold, T. R., Leda, M., Goryachev, A. B., & Miller, A. L. (2019). Rho flares repair local tight junction leaks. Developmental cell, 48(4), 445-459.
6. Yamamoto, K., Miura, H., Ishida, M., Mii, Y., Kinoshita, N., Takada, S., ... & Aoki, K. (2021). Optogenetic relaxation of actomyosin contractility uncovers mechanistic roles of cortical tension during cytokinesis. Nature communications, 12(1), 7145.
7. Thielicke, W., & Sonntag, R. (2021). Particle Image Velocimetry for MATLAB: Accuracy and enhanced algorithms in PIVlab. Journal of Open Research Software, 9(1), 12-12.
8. Zsolnay, V., Gardel, M. L., Kovar, D. R., Voth, G. A. (2024). Cracked actin filaments as mechanosensitive receptors. Biophysical Journal 123(19), 3283–3294.
9. Dominguez, C., Boelens, R., Bonvin, A. M. J. J. (2003). HADDOCK: a protein–protein docking approach based on biochemical or biophysical information. J. Am. Chem. Soc. 125, 1731–1737. (10.1021/ja026939x)
10. van Zundert, G. C. P., Rodrigues, J. P. G. L. M., Trellet, M., Schmitz, C., Kastritis, P. L., Karaca, E., Melquiond, A. S. J., van Dijk, M., de Vries, S. J., Bonvin, A. M. J. J. (2016). The HADDOCK2.2 web server: user-friendly integrative modeling of biomolecular complexes. J. Mol. Biol. 428(4), 720–725. (10.1016/j.jmb.2015.09.014).
11. Weng, G., Wang, E., Wang, Z., Liu, H., Zhu, F., Li, D., Hou, T. (2019). HawkDock: a web server to predict and analyze the protein–protein complex based on computational docking and MM/GBSA. Nucleic Acids Res. 47(W1), W322–W330. (10.1093/nar/gkz397).
12. Massova, I., Kollman, P. A. (2000). Combined molecular mechanical and continuum solvent approach (MM-PBSA/GBSA) to predict ligand binding. Perspect. Drug Discov. Des. 18, 113–135. (10.1023/A:1008763014207)
